# Post-transcriptionally regulated genes are essential for pluripotent stem cell survival

**DOI:** 10.1101/2021.10.18.462967

**Authors:** Mio Iwasaki, Yuka Kawahara, Tsuyoshi Tabata, Yohei Nishi, Megumi Narita, Akira Ohta, Hirohide Saito, Takuya Yamamoto, Masato Nakagawa, Shinya Yamanaka, Kazutoshi Takahashi

## Abstract

The effects of transcription factors on the maintenance and differentiation of pluripotent stem cells (PSCs) have been well studied. However, the importance of post-transcriptional regulatory mechanisms, which cause the quantitative dissociation of mRNA and protein expression, has not been explored in detail. Here, by combining transcriptome and proteome profiling, we identified 228 post-transcriptionally regulated genes with strict upregulation of the protein level in PSCs. Among them, we found that 84 genes were vital for the survival of PSCs and HDFs, including 20 genes that were specifically necessary for the survival of PSCs. These 20 proteins were upregulated only in PSCs and not in differentiated cells derived from the three germ layers. Subcellular fractionation of the mRNA showed that the expression of most of the 20 proteins was regulated at the mRNA localization stage from the nucleus to the cytoplasm, but their translation efficiency was constant. Together, these results revealed that post-transcriptionally regulated genes have a crucial role in PSC survival.

## INTRODUCTION

The processing (Bentley, 2014), export (Hentze et al., 2018), turnover (Dendooven et al., 2020), and accurate decoding of mRNA (Teixeira and Lehmann, 2019), ribosome biogenesis (Pelletier et al., 2018), and protein degradation (Pohl and Dikic, 2019) are critical steps in the post-transcriptional regulation of gene expressions. Among the several hundred human cell types, pluripotent stem cells (PSCs) have unique post-transcriptional mechanisms (Chen and Hu, 2017), such as specific mRNA processing events, that establish and maintain the pluripotent state (Gabut et al., 2011; Han et al., 2013; Ohta et al., 2013; Salomonis et al., 2010). Interestingly, for many genes, differences in protein levels in PSCs are not accompanied by differences in the corresponding mRNA levels (Lu et al., 2009), suggesting post-transcriptionally regulated genes exist in strict regulation at the protein level. One example is the CLOCK gene, in which the amount of mRNA is constant before and after differentiation, whereas the protein expression depends on the cell type and is absent in PSCs (Umemura et al., 2017). Considering that protein levels are more conserved than mRNA levels in primates (Khan et al., 2013), the strict regulation at the protein level with constant mRNA quantities might have important effects on pluripotency and differentiation during development.

To systematically identify genes differently regulated at the mRNA and protein level, transcriptome analysis by RNA sequencing or microarray and proteome analysis by mass spectrometry (MS) have been performed on various cell types (Buccitelli and Selbach, 2020; Matsumoto et al., 2017; Roumeliotis et al., 2017; Wang et al., 2019). Typically, a high correlation in the levels is shown for core metabolic pathway-related genes, but a low correlation is observed for ribosomal and spliceosome genes. However, whether post-transcriptionally regulated genes with tightly controlled protein levels are essential for the maintenance of the cell remains an open question. Here, we compared global mRNA and protein levels between PSCs and differentiated cells to identify essential post-transcriptionally regulated genes. We found 228 post-transcriptionally regulated genes exclusively in PSCs that showed specific biological functions for RNA and nucleic acid binding. siRNA screening revealed that 54% (84 genes) were necessary for cell survival and 20 of them were identified as PSC-specific. Finally, we showed that these 20 genes are mainly regulated at the mRNA localization stage rather than the translation stage, suggesting an importance of cell type-specific post-transcriptional regulation in PSCs.

## RESULTS

### 228 upregulated proteins in PSCs showed constant mRNA expression

First, to identify upregulated genes and post-transcriptionally regulated genes, we analyzed the mRNA and protein levels by microarray and MS, respectively (Figure 1A). We used two human dermal fibroblast (HDF) lines (HDF-1, HDF1388; and HDF-2, Tig120), two human induced PSC (iPSC) lines (iPSC-1, 201B7; and iPSC-2, 1418E1), and one human embryonic stem cell line (ESC, H9). We compared the ratio of the mRNA and protein levels for 6404 genes for all combinations of PSCs (iPSC-1, −2, ESC) and HDFs (HDF-1, −2) (Figure S1). In the six pairs of different cell types, commonly upregulated genes were defined as having at least a 1.9-fold change (log_2_ (0.9)). We found that 1275 genes had upregulated mRNA levels in all PSCs, and 1468 genes had upregulated protein levels (Figure 1B). Among these genes, 509 showed upregulated levels for both mRNA and protein, including PSC-specific genes, such as LIN28A, OCT3/4, and SOX2 (Figure 1B, C). Regarding HDFs, the numbers were 1519 and 461, respectively, and the number with common upregulation was 387, including HDF-specific genes, such as FN1 and VIM (Figure 1B, C). Overall, each cell-specific gene showed a highly correlated mRNA to protein ratio (Figure 1C and Table S1).

**Figure 1.**
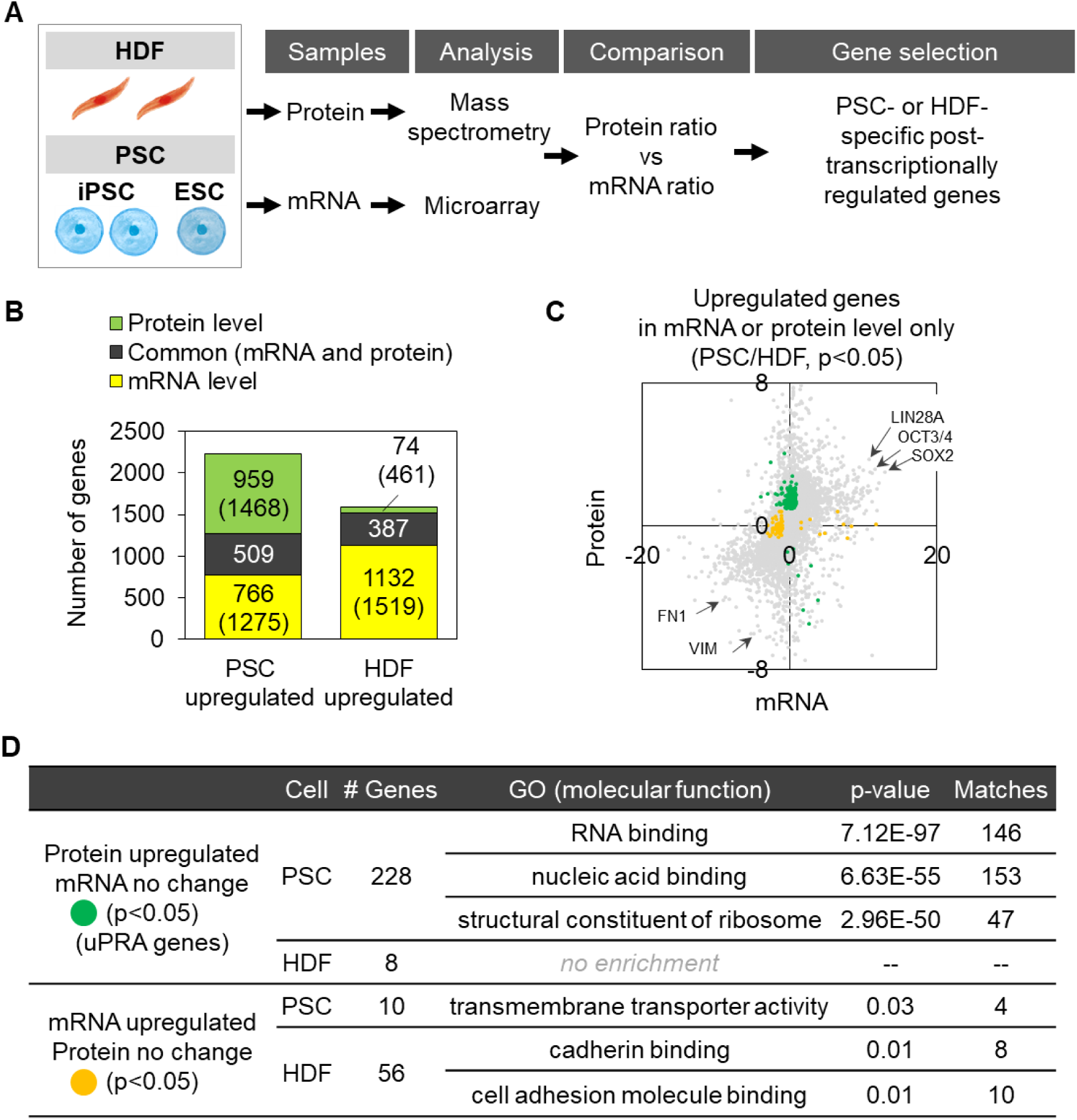
228 and 8 genes were respectively identified as post-transcriptionally regulated genes with independent protein upregulation in PSCs and HDFs. A. Workflow for identifying post-transcriptionally regulated genes using mass spectrometry (MS) for proteins and microarray for mRNA. Two iPSC lines (iPSC-1, 201B7; and iPSC-2, 1418E1), one ESC line (H9), and two HDF lines (HDF-1, HDF1388; and HDF-2, Tig120) were used. B. Number of commonly upregulated genes by protein levels only, mRNA levels only, and both levels in PSCs and HDFs. Numbers in parentheses show the total numbers. C. The mRNA to protein expression ratio in PSCs and HDFs for each gene. Gray, yellow, and green indicate all genes, those with only upregulated mRNA, and those with only upregulated protein levels, respectively (p < 0.05, two-sample unpaired t test). Log_2_ scale. Comparisons of the mRNA and protein ratios between all PSC and HDF lines are shown in Figure S1. The total number of genes quantified was 6404. All plotted data are shown in Table S1. See “Trans-omics data analysis” in Methods for details. D. Number of genes and the GO analysis of molecular functions for genes that were significantly upregulated only at the protein or mRNA level in C.

Next, we examined genes that were post-transcriptionally regulated by selecting those that were upregulated only at the mRNA or protein level by log_2_ (0.9) (Figure 1C; see Trans-omics data analysis in Methods). From this analysis, we found 10 and 56 genes in PSCs and HDFs, respectively, upregulated only at the mRNA level, and 228 and 8 genes upregulated only at the protein level. To investigate the function of these four gene sets, we conducted a Gene Ontology (GO) analysis (Fig. 1D). Especially among the 228 genes in PSCs that showed only upregulated protein levels, we found specific molecular functions, such as RNA binding and nucleic acid binding. Although the 8 genes for HDFs showed no enrichment in molecular functions, we named these 228 and 8 genes “<u>u</u>pregulated <u>p</u>rotein levels independent of m<u>R</u>N<u>A</u> levels (uPRA)” for further analysis and found that 228 PSC-uPRA genes have specific nucleic acid binding functions.

### 20 uPRA genes are essential for PSC cell survival

To investigate which uPRA genes are significant for cell maintenance, we performed siRNA knockdown screening in iPSC-1, −2, and HDF-2 (Figure 2A, 2B, Table S2, S3). We picked siRNAs to target 156 (Table S2) of the 228 PSC-uPRA and 8 HDF-uPRA genes. The siRNA knockdown of OCT3/4 and LaminB2 was respectively used as a pluripotency marker for PSCs and a universal control for both cell types (Figure 2B). The knockdown of none of the 156 uPRA genes induced differentiation (Table S3). On the other hand, the knockdown of 84 of the 156 uPRA genes, such as SRRT and RSL1D1, significantly decreased the number of PSCs (Figure 2B). Among them, the knockdown of 52 uPRA genes decreased the number of HDFs too (Figure 2C). This result indicates that uPRA genes have important effects on the cellular maintenance of HDFs as well. Additionally, we identified 20 PSC-uPRA genes and 1 HDF-uPRA gene that are each specifically necessary for the survival of the respective cells (Figure 2C; images of the knockdown of these 21 genes are shown in Figure S2). A GO analysis showed that the 20 PSC-uPRA genes code for components of ribonucleoprotein and protein-containing complexes and have RNA- and nucleic acid-binding properties (Figure 2D, E). These data suggest that the 20 PSC-uPRA genes are essential for the survival of PSCs and synergistically maintain PSCs via heterocyclic compound binding properties.

**Figure 2.**
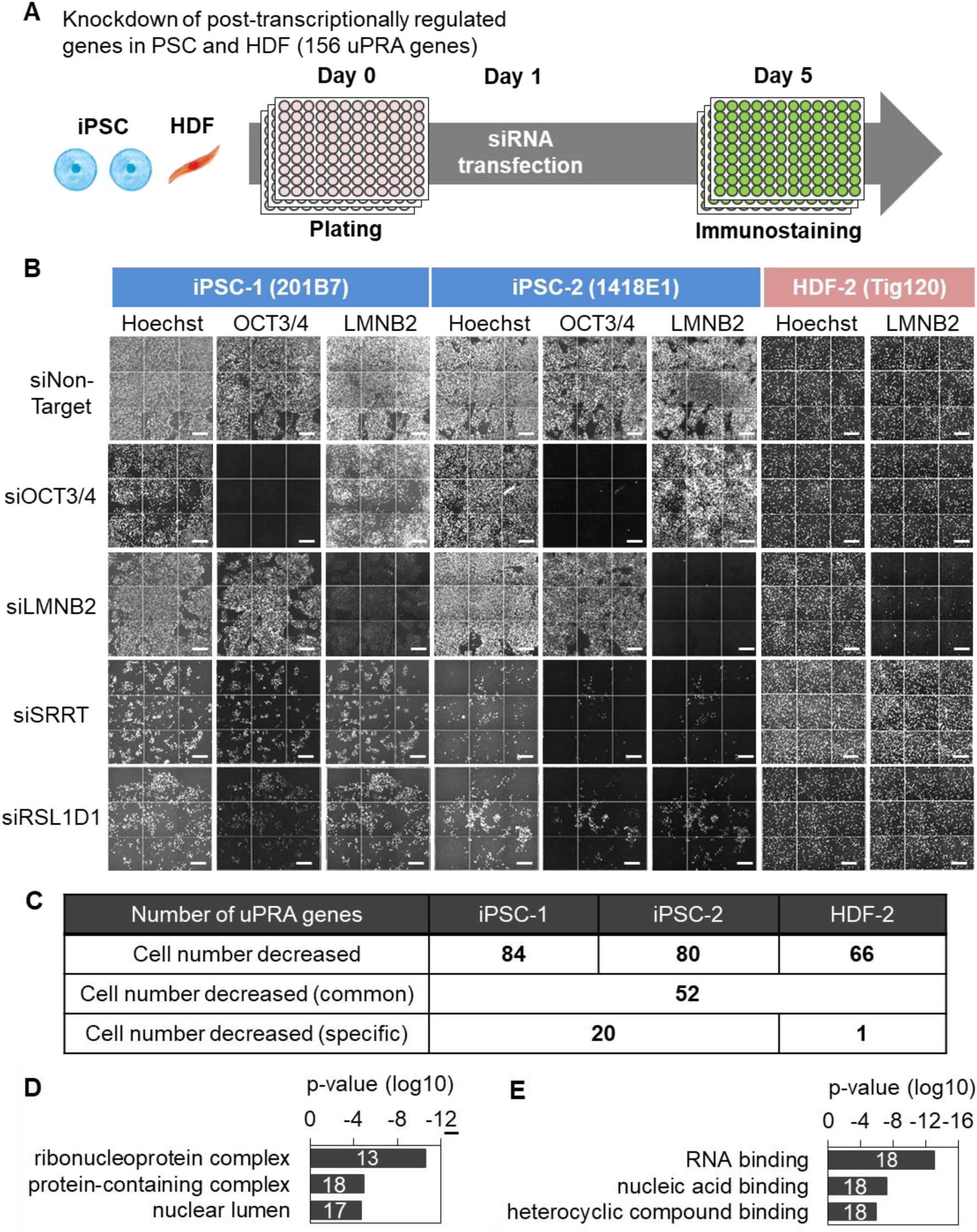
20 uPRA genes in PSCs and 1 uPRA gene in HDFs were related to cell survival. A. Workflow of the knockdown experiment by siRNA for uPRA genes in PSCs and HDFs (156 uPRA genes in total). iPSC-1 (201B7), iPSC-2 (1418E1), and HDF-2 (Tig120) were used for the assay. See “siRNA screening” in Methods for details. The knockdown targets are shown in Table S2, and the immunoassay cell percentage and observed number of cells are shown in Table S3. B. Representative images of the siRNA knockdown experiment. Nuclei were visualized by Hoechst 33342, and PSCs were visualized by OCT3/4 expression. The knockdown efficiency was visualized by OCT3/4 and LMNB2 expression. siSRRT and siRSL1D1 are representative images for siRNAs experiments in which the cell number decreased. Bars indicate 200 μm. C. The number of post-transcriptionally regulated genes (uPRA genes) whose knockdown caused a substantial decrease in cell number. Images for the siRNAs of the total 21 cell-specific uPRA genes are shown in Figure S2. D. GO analysis of cellular component for the 20 PSC-uPRA genes in C. E. GO analysis of molecular function for the 20 PSC-uPRA genes in C.

### Upregulated protein levels of the 20 essential uPRA genes are PSC-specific

We examined if the 20 PSC-uPRA genes are regulated post-transcriptionally only in PSCs. We selected another seven differentiated primary cell lines representing the three germ layers to compare with the PSCs, which are undifferentiated. In total, mesoderm-derived cells included adipose tissue-derived mesenchymal stem cells (HAdMSC) and two HDF lines (HDF-1 and 2), endoderm-derived cells included normal human bronchial epithelial cells (NHBEC) and human prostate epithelial cells (PrEC), and ectoderm-derived cells included normal human epidermal keratinocytes (NHEK) and human neural progenitor cells derived from H9 (NPC H9). Quantitative reverse transcription PCR (qRT-PCR) and immunoblotting were performed on the mRNA and proteins, respectively, and the gene expressions relative to HDF-1 were compared (all blotting images and graphs are shown in Figure S3A, C). RSL1D1, one of the 20 essential PSC-uPRA genes, showed a more than 50-fold increase in its protein expression only in PSCs, whereas its mRNA expression was equal among all cell types (Figure 3A, B). Overall, we confirmed that the protein expressions of 18 PSC-uPRA genes (RBM22 and SF3B3 were the exceptions) were increased more than two-fold compared with their mRNA expressions in PSCs (Figure 3C). On the other hand, the fold-change between protein and mRNA expressions was around one or less in differentiated cells for 17 PSC-uRPO genes (IMP4, NCBP2, and BUD31 were the exceptions), suggesting the post-transcriptional regulation mechanism is different between PSCs and differentiated cells (Figure 3D). RSL1D1, SF3B4, RBM22, and SF3B3 especially showed protein to mRNA expression ratios less than one in differentiated cells, indicating that the protein expressions of these genes are usually suppressed in cell types other than PSCs. As for AFP, the one HDF-uPRA gene, we confirmed a specific increase in the protein expression in mesoderm-derived cells (Figure S3B). Following these observations, we concluded that the 20 PSC-uPRA genes are post-transcriptionally regulated specifically in PSCs.

**Figure 3.**
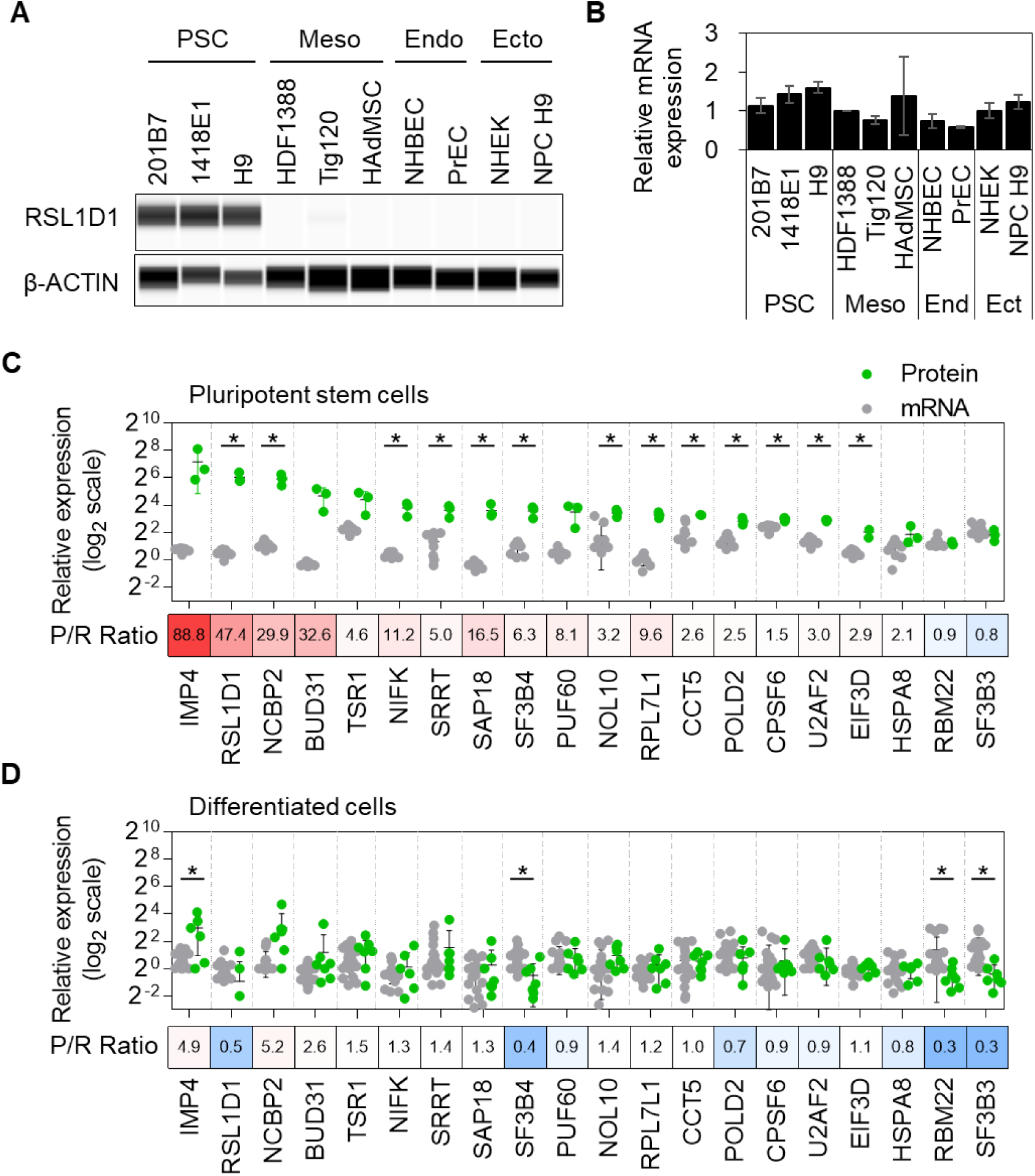
20 uPRA genes were post-transcriptionally regulated only in PSCs. A. A representative image of the immunoblotting for one PSC-uPRA gene, RSL1D1, in various cell types. We used three cell lines for PSCs, three cell lines for mesoderm-derived cells, two cell lines for endoderm-derived cells, and two cell lines for ectoderm-derived cells for the gene expression analysis. Immunoblotting images of all 20 essential PSC-uPRA genes are shown in Figure S3A. B. Gene expression measured by qRT-PCR was normalized by GAPDH. The relative expression ratios were normalized to the result of HDF1388 (HDF-1). Representative results for the RSL1D1 mRNA are shown as the mean ± S.D. Biological triplicates for the mRNA quantification. Results for all essential PSC-uPRA genes are shown in Figure S3C. C. Relative gene expressions of mRNA and protein levels in PSCs for the 20 essential PSC-uPRA genes normalized to HDF1388 (HDF-1). A heatmap of the protein to mRNA expression ratio (P/R ratio) is shown above each gene name. *p < 0.05, two-sample unpaired t test. D. Relative mRNA and protein levels in differentiated cells for the 20 essential PSC-uPRA genes normalized to HDF1388 (HDF-1). A heatmap of the P/R ratio is shown above each gene name. *p < 0.05, two-sample unpaired t test.

### HSPA8, EIF3D, and NCBP2 protein expressions are controlled at the ubiquitin-dependent degradation stage

Next, we analyzed individual post-transcriptional regulation stages to identify the regulation mechanism of the protein expressions of the 20 essential PSC-uPRA genes (Figure 4A). After transcription, mRNAs are transported from the nucleus to the cytoplasm, where ribosomes bind to translate them. Eventually, the proteins are degraded by the ubiquitin-proteasome system and/or lysosomes. At first, we analyzed the protein degradation efficiency using MG-132 (proteasome inhibitor) and Bafilomycin A1 and Wortmannin (lysosome inhibitors) as controls for the assays (Figure S4A-D). These inhibitors should recover the PSC-uPRA protein expression in HDFs if the proteins are downregulated by fast degradation. We found that the protein expressions of HSPA8, EIF3D, NCBP2, and IMP4 were steadily increased more than two-fold after exposure to proteasome inhibitor for up to 8 hours (Figure 4B, S4E). We did the same assay for up to 24 hours and confirmed that the protein expressions of HSPA8, EIF3D, and NCBP2 were controlled by fast degradation via the proteasome system in HDFs (Figure 4C). The lysosome inhibitor assay (Figure 4D, S4F) showed high protein expressions of NCBP2 and RSL1D1 for treatment up to 8 hours but not 24 hours (Figure 4E), suggesting no PSC-uPRA genes were regulated by lysosome-dependent degradation. Overall, these data indicate that the protein expressions of three PSC-uPRA genes, HSPA8, EIF3D and NCBP2, are regulated at the ubiquitin dependent degradation stage.

**Figure 4.**
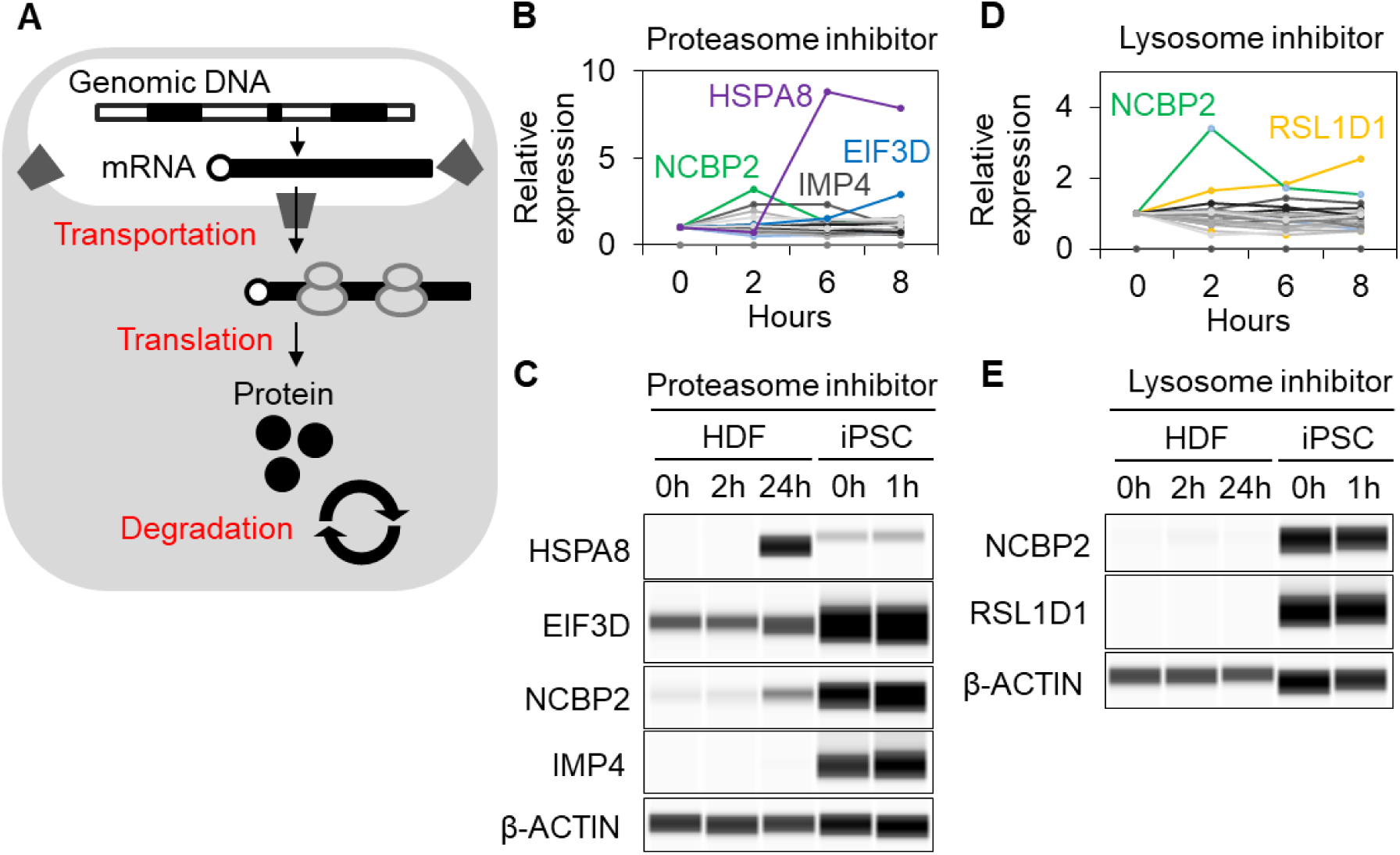
HSPA8, EIF3D, and NCBP2 protein expressions are controlled at the ubiquitin-dependent degradation stage. A. Overview of the known regulation stages for gene expression after transcription. B. HDF-2 (Tig120) was treated with a proteasome inhibitor (20 μM MG-132) for up to 8 hours, and the expression levels of the 20 essential PSC-uPRA genes were measured by immunoblotting. The relative expression of each protein compared to pre-inhibition is shown. All immunoblotting images are shown in Figure S4A. C. HDF-2 (Tig120) and iPSC-1 (201B7) were treated with 20 μM MG-132 for up to 24 hours, and the protein expressions for HSPA8, EIF3D, NCBP2, and IMP4 were measured by immunoblotting. D. HDF-2 (Tig120) was treated with lysosome inhibitors (250 nM Bafilomycin A1 and 500 nM Wortmannin) for up to 8 hours, and the protein expression of the 20 essential PSC-uPRA genes were measured by immunoblotting. The relative expression of each protein to pre-inhibition is shown. All immunoblotting images are shown in Figure S4B. E. HDF-2 (Tig120) and iPSC-1 (201B7) were treated with 250 nM Bafilomycin and 500 nM Wortmannin for up to 24 hours, and the protein expressions of NCBP2 and RSL1D1 were measured by immunoblotting.

### The cytosolic localization but not the translation efficiency of uPRA mRNA is increased in PSCs

Next, we analyzed the subcellular localization of the mRNAs of the 20 essential PSC-uPRA genes to examine their intracellular abundance between different cell types. We used 18S rRNA and MALAT1 lncRNA, respectively, as cytosolic and nuclear RNA controls in the subcellular fractionation experiment (Figure 5A). For cytosolic RNA, 94.3% and 73.3% of 18S rRNA were observed in PSCs and HDFs, respectively, and for nuclear RNA, 99.5% and 99.8% of MALAT1 lncRNA were observed, validating the subcellular fractionation experiment. Then we analyzed the cytosolic RNA percentage of the 20 essential PSC-uPRA genes by qRT-PCR (Figure 5B). The cytosolic RNA percentage of 18 PSC-uPRA genes (IMP4 and CPSF6 were the exceptions) were increased at least more than two-fold in PSCs. Conversely, the nuclear RNA percentage of PSC-uPRA genes was increased in HDFs (Figure 5C). These results indicated that the mRNAs of the 18 PSC-uPRA genes were retained in the nucleus in HDFs.

**Figure 5.**
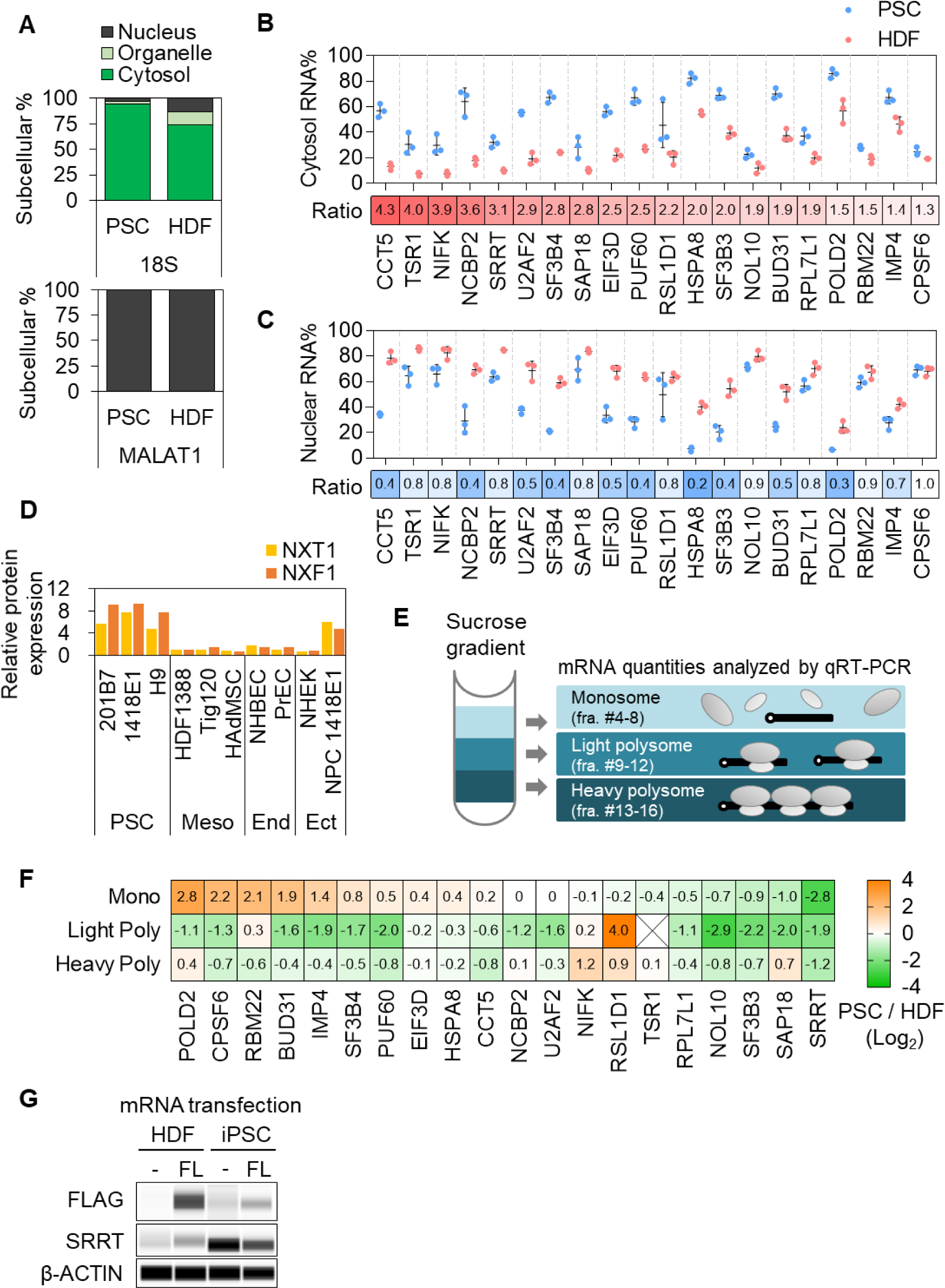
PSC-uPRA mRNAs are retained in the nucleus of HDFs without changing the translation efficiency. A. The percentages of mRNA in the cytoplasm, membrane, and nucleus of iPSC-1 (201B7) and HDF-2 (Tig120) are shown for 18S ribosomal RNA and MALAT1 lncRNA measured by qRT-PCR. B. The percentage of mRNA in the cytoplasm measured by qRT-PCR for the 20 essential PSC-uPRA genes. The cytoplasm mRNA expression ratio in PSCs to HDFs is shown above each gene name in the heatmap. C. The percentage of mRNA in the nucleus measured by qRT-PCR for the 20 essential PSC-uPRA genes. The nucleus mRNA expression ratio in PSCs to HDFs is shown above each gene name in the heatmap. D. The protein-fold change between PSCs and other differentiated cells for NXT1 and NXF1. Protein expressions measured by immunoblotting were normalized by b-ACTIN. The relative expression ratios were normalized to the result of HDF1388 (HDF-1). E. Overview of the analysis of the translation efficiency by the sucrose gradient. The monosome (fraction 4–8), light polysome (fraction 9–12), and heavy polysome (fraction 13–16) were collected to extract RNAs, which were measured by qRT-PCR. See “Monosome and polysome fractionation” in Methods for details. F. The percentage of mRNAs in monosome (Mono), light polysome (Light Poly), and heavy polysome (Heavy Poly) were analyzed by qRT-PCR. The average mRNA expression ratio in PSCs to HDFs is shown in the heatmap with log_2_ scale (n=3). Values are normalized by spike RNA and compared with the loading sample before the sucrose gradient. G. Full-length FLAG-SRRT mRNA was transfected into HDF-2 (Tig120) and iPSC-1 (201B7). Immunoblotting assays for FLAG, SRRT, and b-ACTIN are shown.

RNA splicing is known to promote mRNA-export to the cytoplasm, because export factors can be recruited by the spliceosome to mature RNA (Palazzo and Lee, 2018). However, PSCs and differentiated cells do not express different spliced isoforms of the 20 PSC-uPRA genes (Han *et al*., 2013). We then checked the protein expressions of RNA export factors. We found more than a four-fold increase in the protein expressions of NXT1 and NXF1, two mRNA export receptor heterodimers, in PSCs and NPC 1418E1 compared to the other differentiated cells (Figure 5D, S5A, B). These data indicated that most PSC-uPRA mRNA was retained in the nucleus of HDFs, but the increased NXT1 and NXF1 protein expressions may accelerate mRNA transport to the cytoplasm in PSCs.

Next, to investigate the translation efficiency, we extracted mRNAs in the monosome, light polysome, and heavy polysome fractions of PSCs and HDFs using sucrose fractionation. We quantified and compared the PSC-uPRA mRNA in each fraction between PSCs and HDFs (Figure 5E). Except for NIFK and RSL1D1, no PSC-uPRA mRNA was enriched in the heavy polysome fraction of PSCs (Figure 5F). This result suggests that the translation efficiency of most PSC-uPRA mRNAs cannot explain the increased protein levels in PSCs.

Among PSC-uPRA mRNAs, SRRT mRNA was the most associated with heavy polysomes in HDFs despite the inhibited protein expression (Figure 5F). Since a lower cytosolic RNA percentage of endogenous SRRT was observed in HDFs (around 10%; Figure 5B), we next analyzed whether its translation occurs if the cytoplasmic RNA content is increased. We constructed full-length FLAG-tagged SRRT mRNA and one day after the mRNA transfection confirmed the translation of FLAG-SRRT protein from the transfected mRNA by immunoblotting (Figure 5G). In summary, these results indicate that there exists an inhibitory mechanism in HDFs for transporting PSC-uPRA mRNAs to the cytoplasm and for the translation of PSC-uPRA genes even after mRNA binds to ribosomes.

## DISCUSSION

Despite their ubiquitous effects, there is comparatively little understanding about post-transcriptional regulatory mechanisms compared with transcriptional regulatory mechanisms in PSCs. Here, we revealed that 20 post-transcriptionally regulated genes (i.e. uPRA genes) are essential for the maintenance of PSCs. These genes had upregulated protein levels without any major positive upregulation in mRNA levels in PSCs (Figure 3C). The regulatory processes of these 20 essential PSC-uPRA genes are illustrated in Figure 6. In HDFs, 18 PSC-uPRA mRNAs were highly retained in the nucleus (Figure 5C). Among them, the translation efficiency of RSL1D1 and NIFK was upregulated in PSCs (Figure 5F), and the proteins of HSPA8, EIF3D, and NCBP2 were quickly degraded in HDFs at the proteasome degradation stage (Figure 4B, C). Also, we found that the translation efficiency of 18 uPRA genes was comparable between PSCs and HDFs by polysome profiling (Figure 5F), suggesting several inhibitory effects for uPRA mRNA transportation to the cytoplasm and for translation after ribosome binding in differentiated cells.

**Figure 6.**
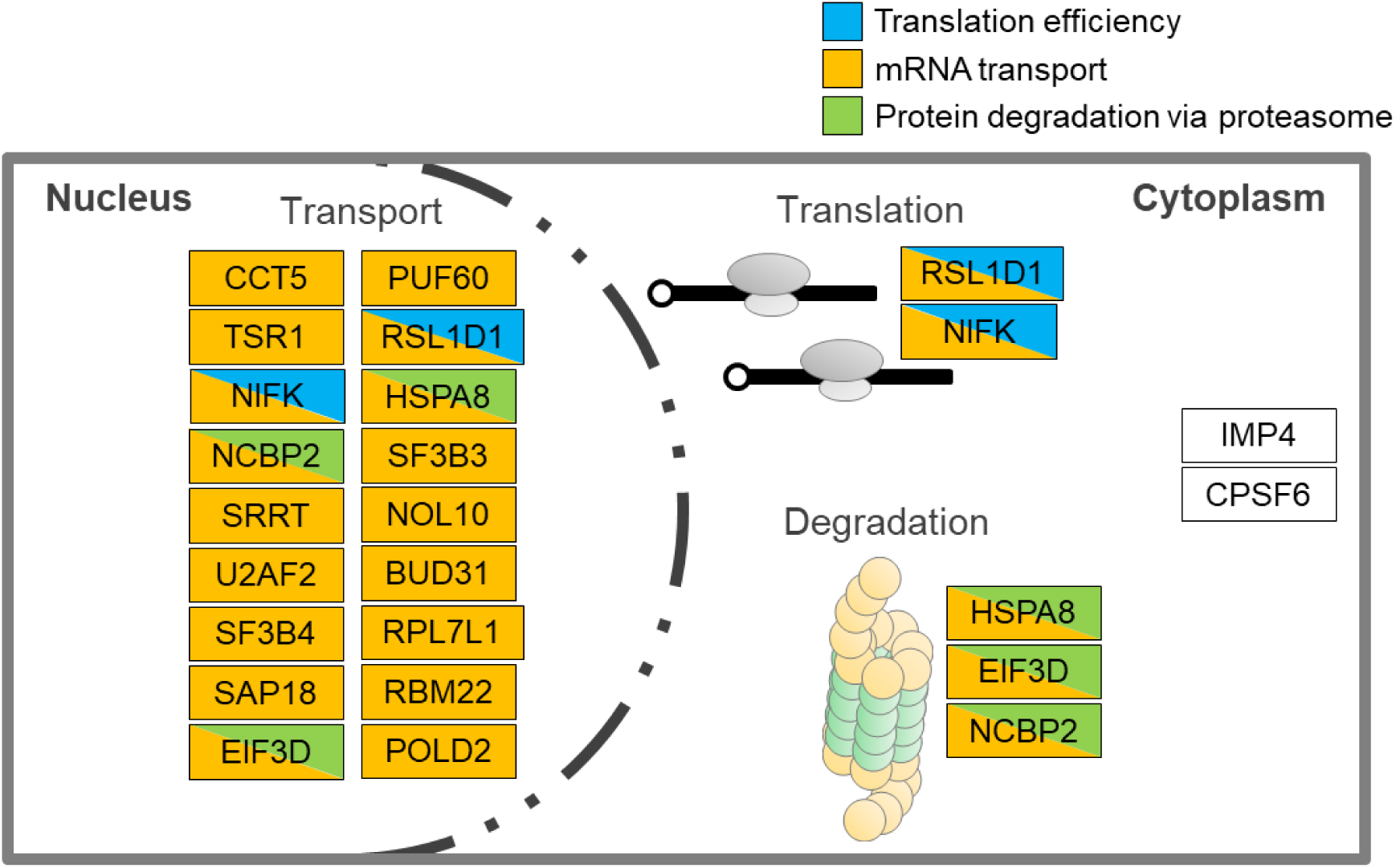
An illustration of how 20 essential PSC-uPRA genes are regulated in PSCs and differentiated cells. Blue, yellow, and green indicate the translation efficiency, mRNA transport, and protein degradation via the proteasome of the 20 essential PSC-uPRA genes, respectively. More mRNA of 18 uPRA genes was transported from the nucleus to the cytoplasm in PSCs than in HDFs. The proteins of HSPA8, EIF3D, and NCBP2 were quickly degraded in HDFs at the proteasome degradation stage. The translation efficiency of RSL1D1 and NIFK was higher in PSCs compared to HDFs. No regulation was observed for IMP4 or CPSF6.

To identify the inhibitory effect, we picked up one uPRA gene, SRRT, for further analysis. The percentage of cytoplasmic SRRT mRNA was 3-fold less in HDFs (Figure 5B), and the mRNA itself was more associated with heavy polysomes in HDFs than in PSCs (Figure 5F). Furthermore, exogenous cytosolic SRRT mRNA was translated in HDFs (Figure 5G), suggesting the cytoplasmic mRNA level is key for proper protein expression. Another possible mechanism for translation inhibition in differentiated cells is ribosome pausing, in which the mRNA binds to ribosomes that are not active or select non-canonical start codons (Chandrasekaran et al., 2019; Darnell et al., 2018; Tresenrider et al., 2021). A third possibility is that aberrant proteins were degraded immediately after the production by a quality control pathway of the mRNA and ribosome (D’Orazio and Green, 2021). More study is needed to clarify which mechanism applies to which gene.

All identified 20 PSC-uPRA proteins in the present study are involved in a wide range of cellular functions, from transcription to post-translation. For transcription-related functions, 10 PSC-uPRA genes have polymerase-, splicing- and mRNA maturation-related functions: POLD2 is a component of the DNA polymerase complex; SF3B3, SF3B4, and RBM22 are components of the splicing factor; and PUF60, BUD31 SAP18, CPSF6, U2AF2, and SRRT are required for the splicing of pre-mRNA, with SRRT especially known as “a molecular guardian of the pluripotent cell state” by facilitating proper splicing in PSCs (Kainov and Makeyev, 2020). For translation-related functions, 8 PSC-uPRA genes are involved in functions of the ribosome complex: NCBP2 and EIF3D are mRNA cap binding proteins; EIF3D is necessary for specialized translation initiation factors (Lee et al., 2016); RPL7L1 is a putative component of the ribosome complex; and RSL1D1, NIFK, NOL10, IMP4, and TSR1 are required for rRNA processing. For post-translational functions, CCT5 and HSPA8 are components of the chaperone complex, which helps newly synthesized proteins properly fold. Our observation that these genes are post-transcriptionally regulated is particularly interesting, because they all function as regulators post-transcription. Thus, they might use feedback in their regulation to maintain pluripotency.

In conclusion, our study revealed that the protein expressions of 228 genes are post-transcriptionally regulated in PSCs. The protein levels of 20 of these genes are specifically increased in PSCs, and the proteins themselves are essential for PSC maintenance. Finally, different mechanisms regulate the gene expressions between cell types.

### Limitation of the study

We focused on post-transcriptionally regulated genes essential for the maintenance of PSCs. Although we showed critical regulatory steps for controlling uPRA protein levels, a wide variety of regulatory mechanisms seem to be involved. Recent studies have shown that the secondary structure of mRNA is important for regulating protein expression by changing the interaction with the expansion segment of ribosomal RNA (Leppek et al., 2020) and changing the mRNA half-life (Mauger et al., 2019). Future studies are needed to determine if secondary structures affect the translation of the 20 PSC-uPRA genes.

## Supporting Information

**Figure S1.**
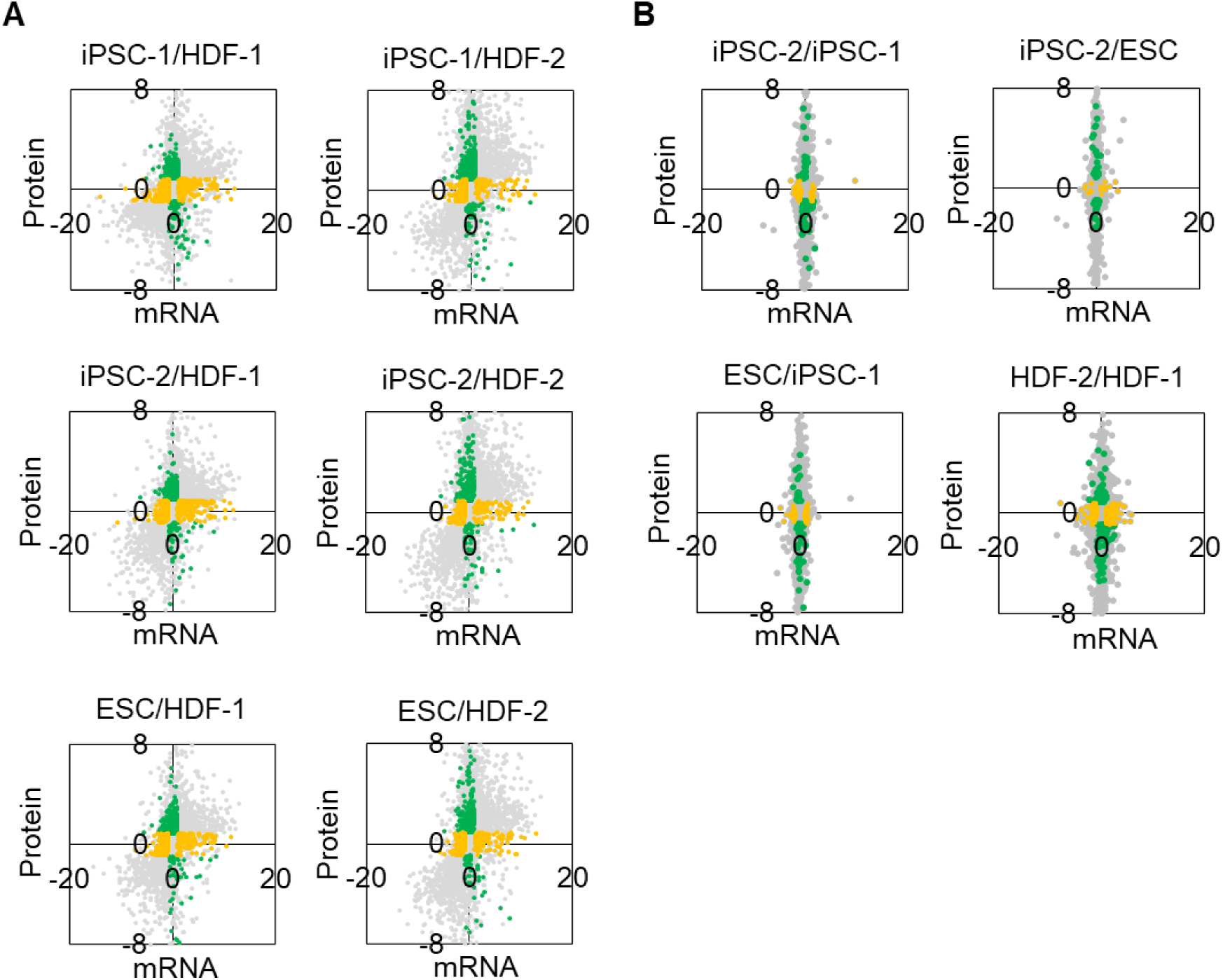
Comparison of the mRNA to protein ratio between cell lines A. The mRNA to protein ratio for each gene between different PSC and HDF lines. Genes with significantly upregulated mRNA and protein levels only are shown in yellow and green, respectively (p < 0.05, two sample paired t test). B. The mRNA to protein ratio for each gene between the same cell type. Genes with significantly upregulated mRNA and protein levels only are shown in yellow and green, respectively (p < 0.05, two sample paired t test).

**Figure S2.**
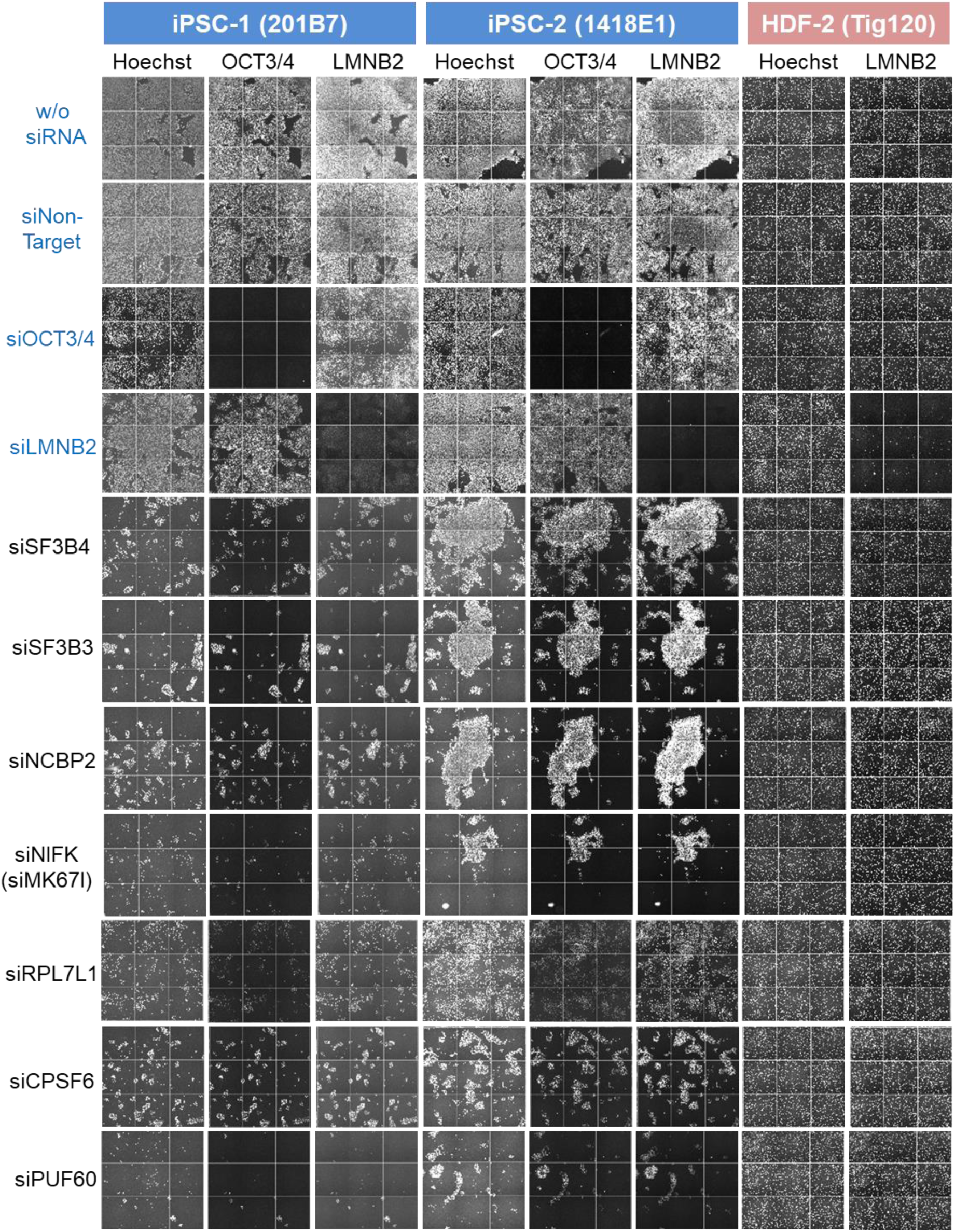
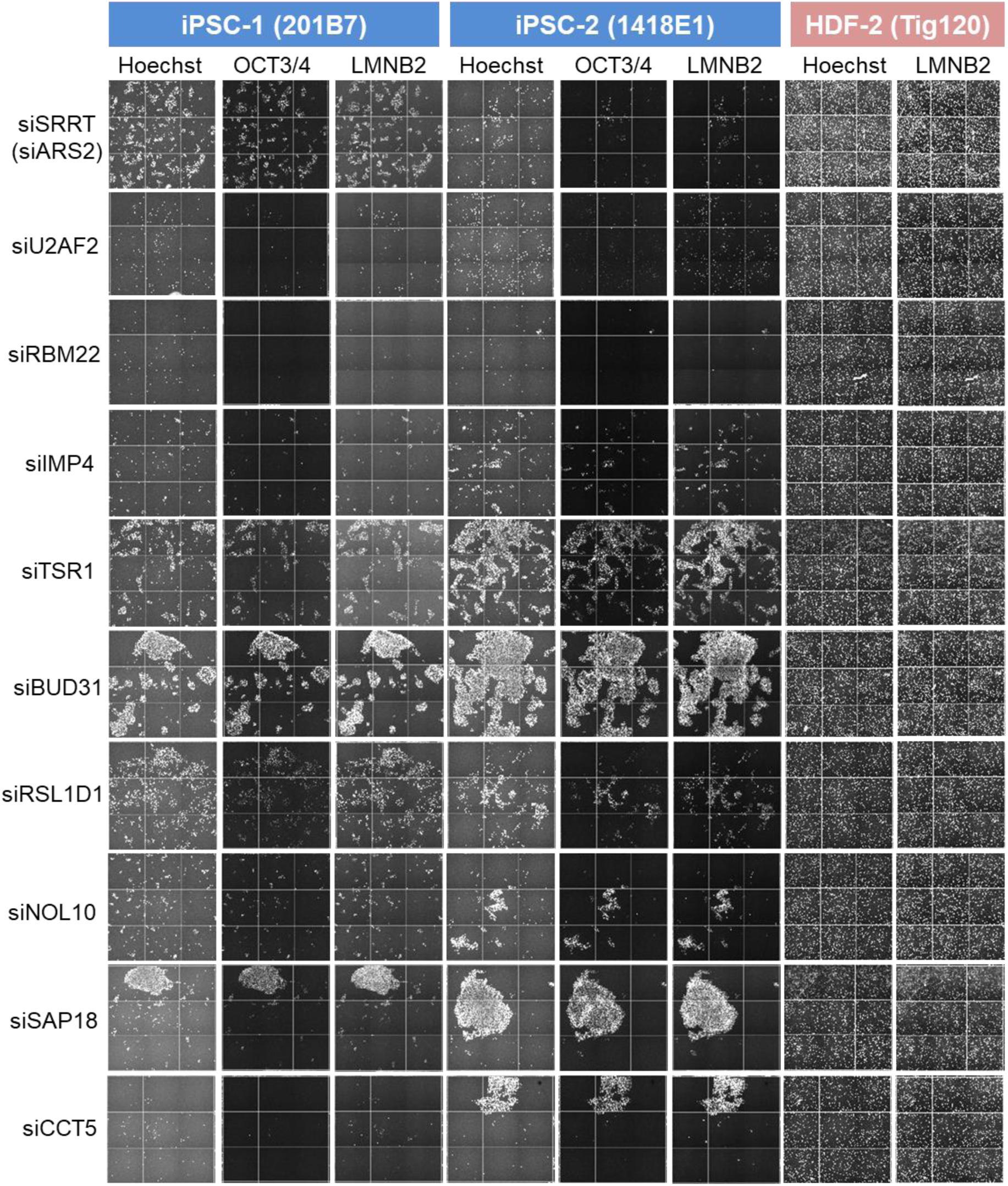
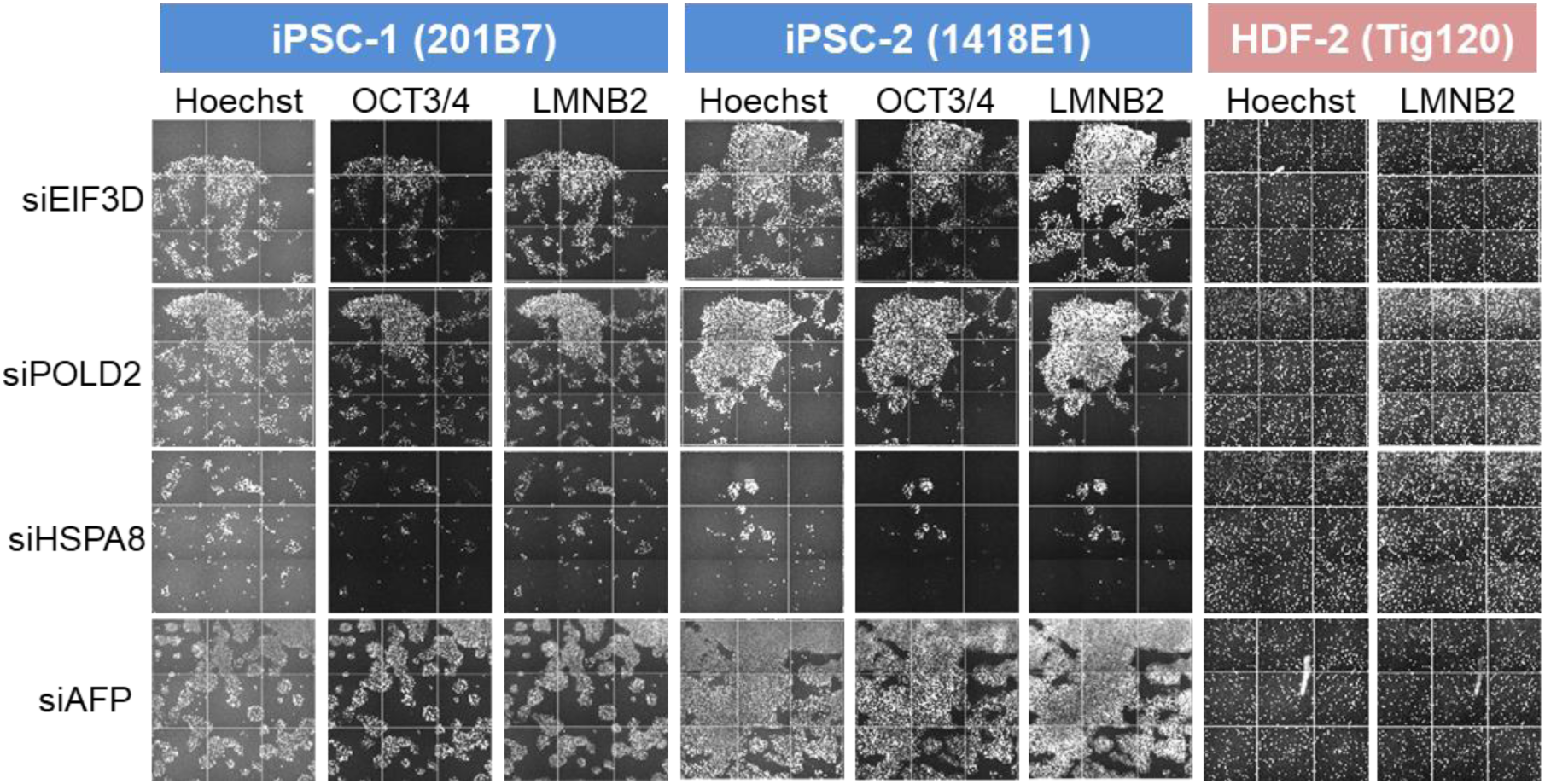
Images of 20 PSC-uPRA genes and 1 HDF-uPRA gene whose knockout caused a decrease in cell survival. Hoechst 33342-stained nuclei, OCT3/4-stained PSCs, and LaminB2 (LMNB2)-stained PSCs and HDFs are shown. Knockdown efficiency was visualized by siRNA for OCT3/4 and LaminB2. The 20 essential PSC-uPRA genes are SF3B4, SF3B3, NCBP2, NIFK, RPL7L1, CPSF6, PUF60, SRRT, U2AF2, RBM22, IMP4, TSR1, BUD31, RSL1D1, NOL10, SAP18, CCT5, EIF3D, POLD2, and HSPA8. The one essential HDF-uPRA gene is AFP.

**Figure S3.**
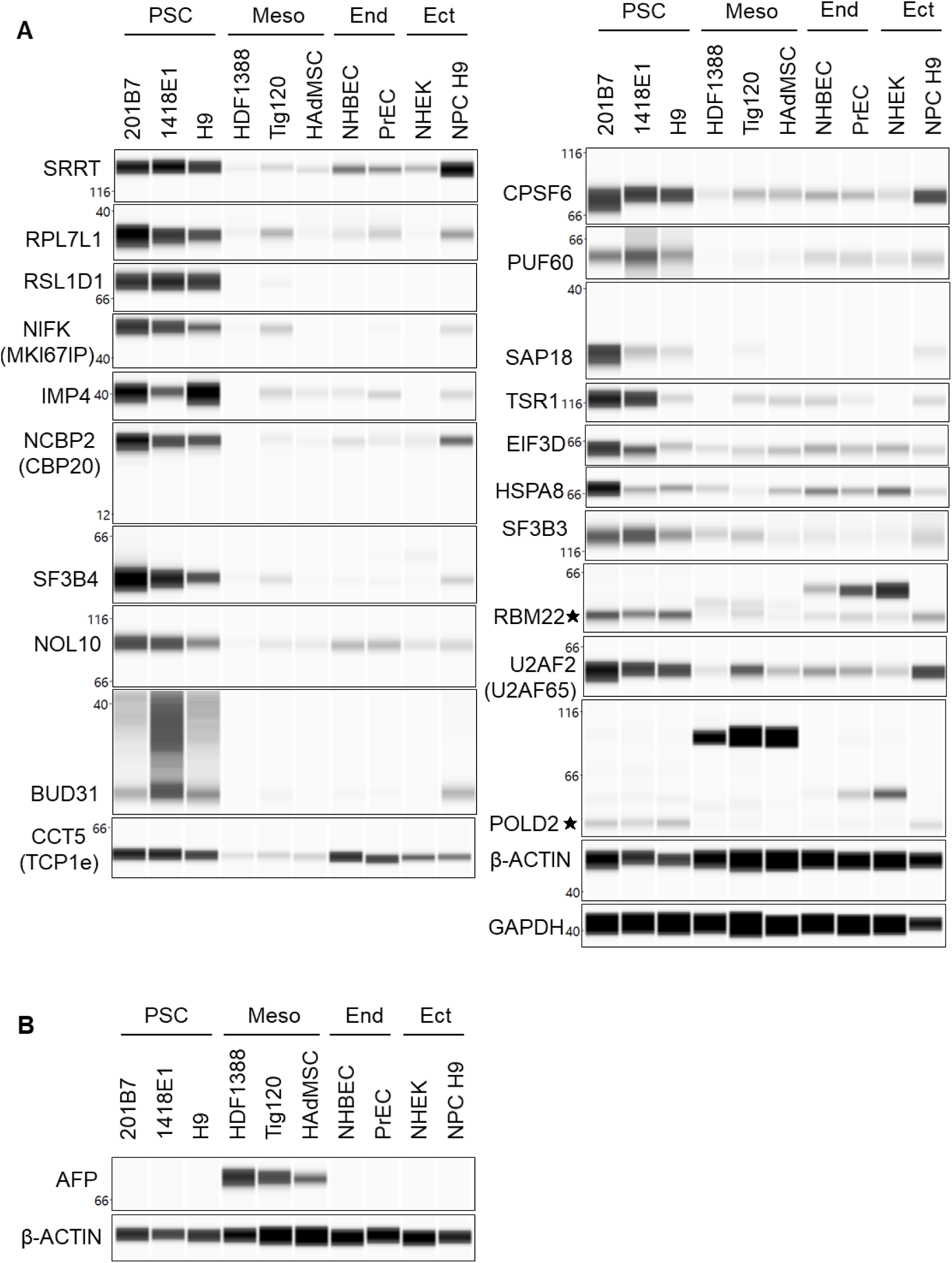
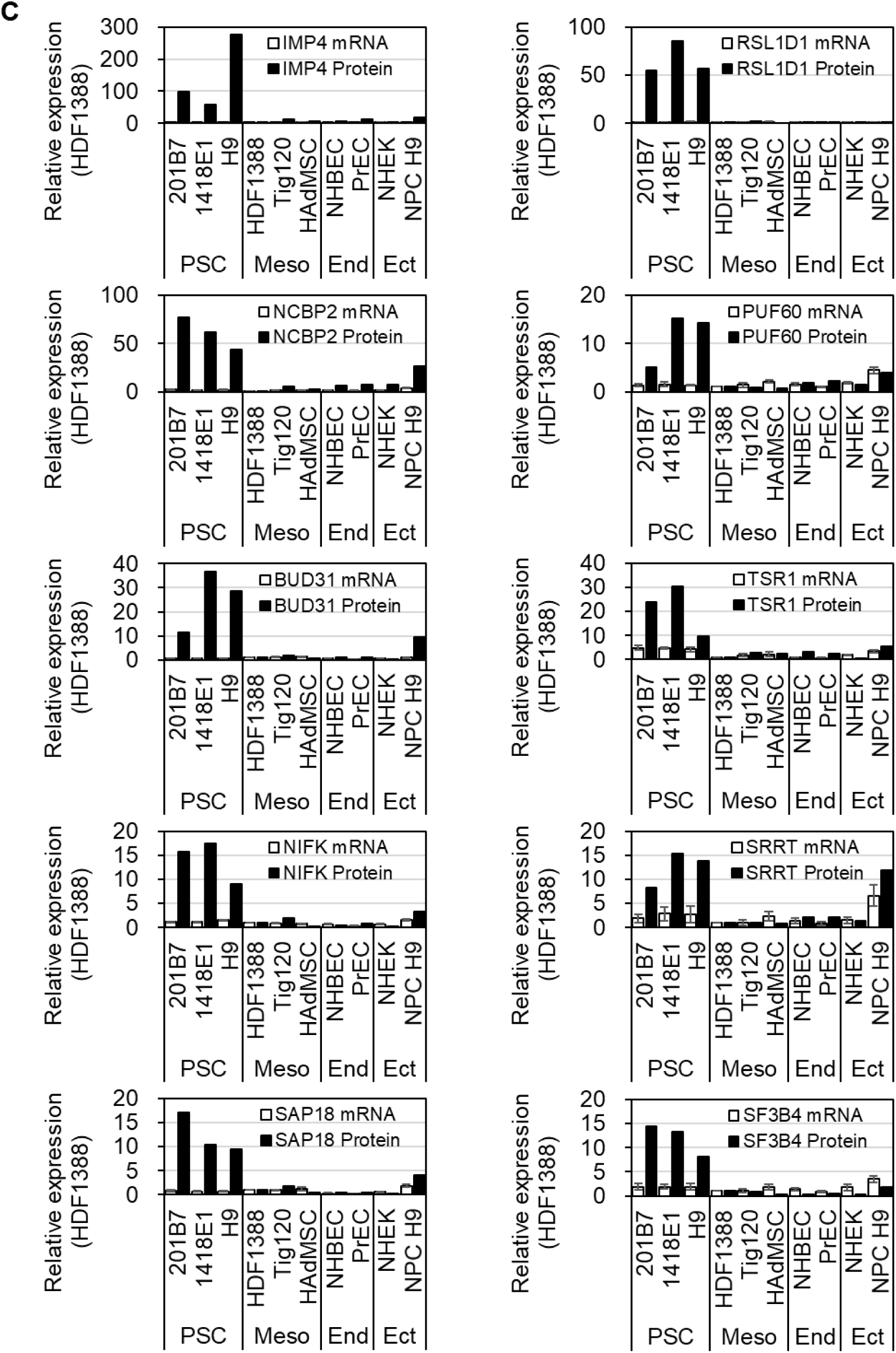
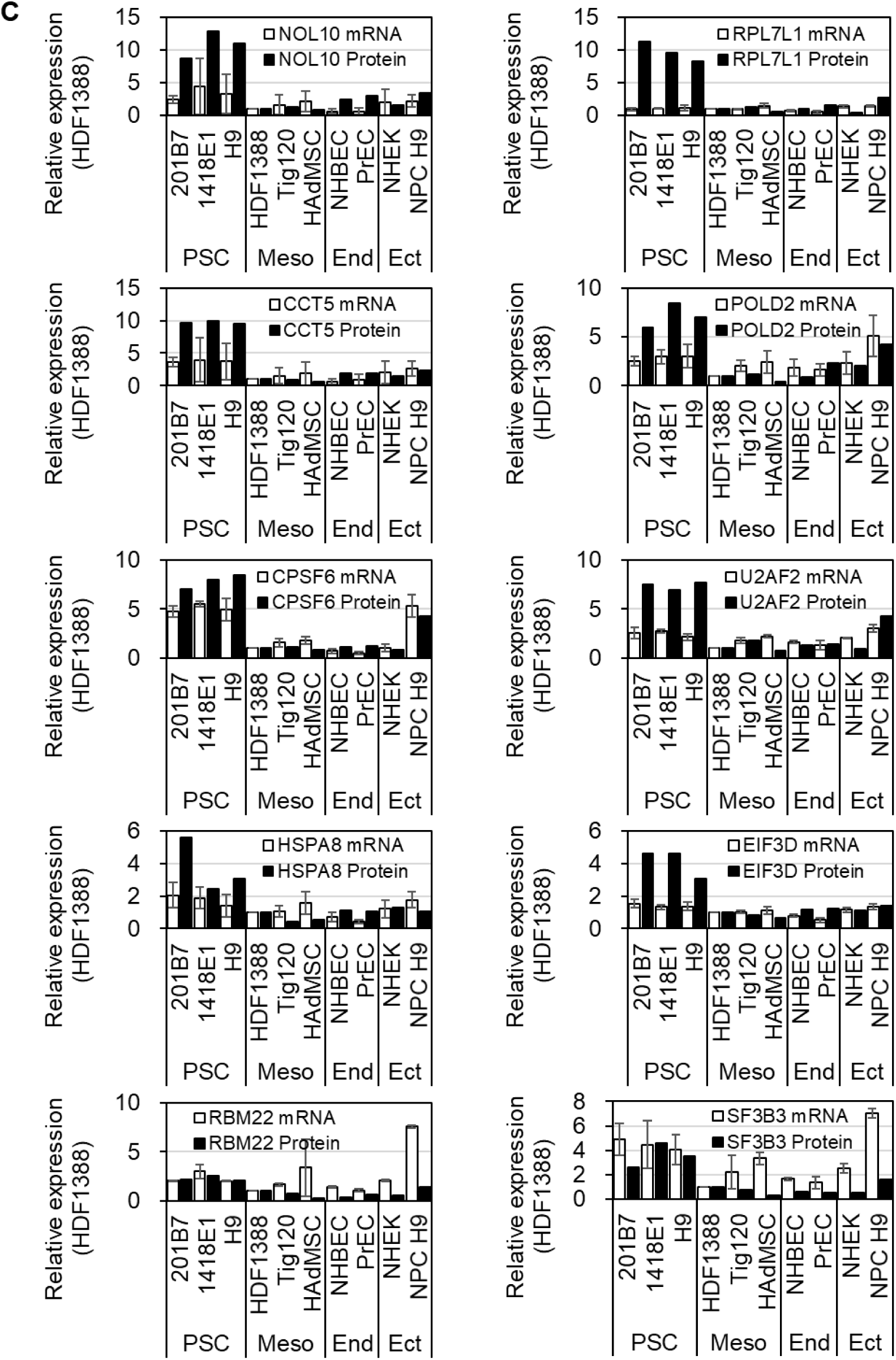
Immunoblotting images and relative expressions of the 20 uPRA genes and one HDF-uPRA gene. A. Immunoblotting images of the 20 essential PSC-uPRA genes. Meso, mesoderm; End, endoderm; Ect, ectoderm. Three cell lines were used for PSCs, three cell lines for mesoderm-derived cells, two cell lines for endoderm-derived cells, and two cell lines for ectoderm-derived cells. B. Immunoblotting image of the 1 essential HDF-uPRA gene (AFP). We could not quantify the mRNA by qRT-PCR because of the low mRNA level. C. Gene expressions measured by qRT-PCR and immunoblotting normalized to HDF-1 (HDF1388). For the mRNA quantification, GAPDH was used for the normalization, and bars represent mean ± S.D., biological triplicate. For the protein quantification, β-ACTIN was used for the normalization.

**Figure S4.**
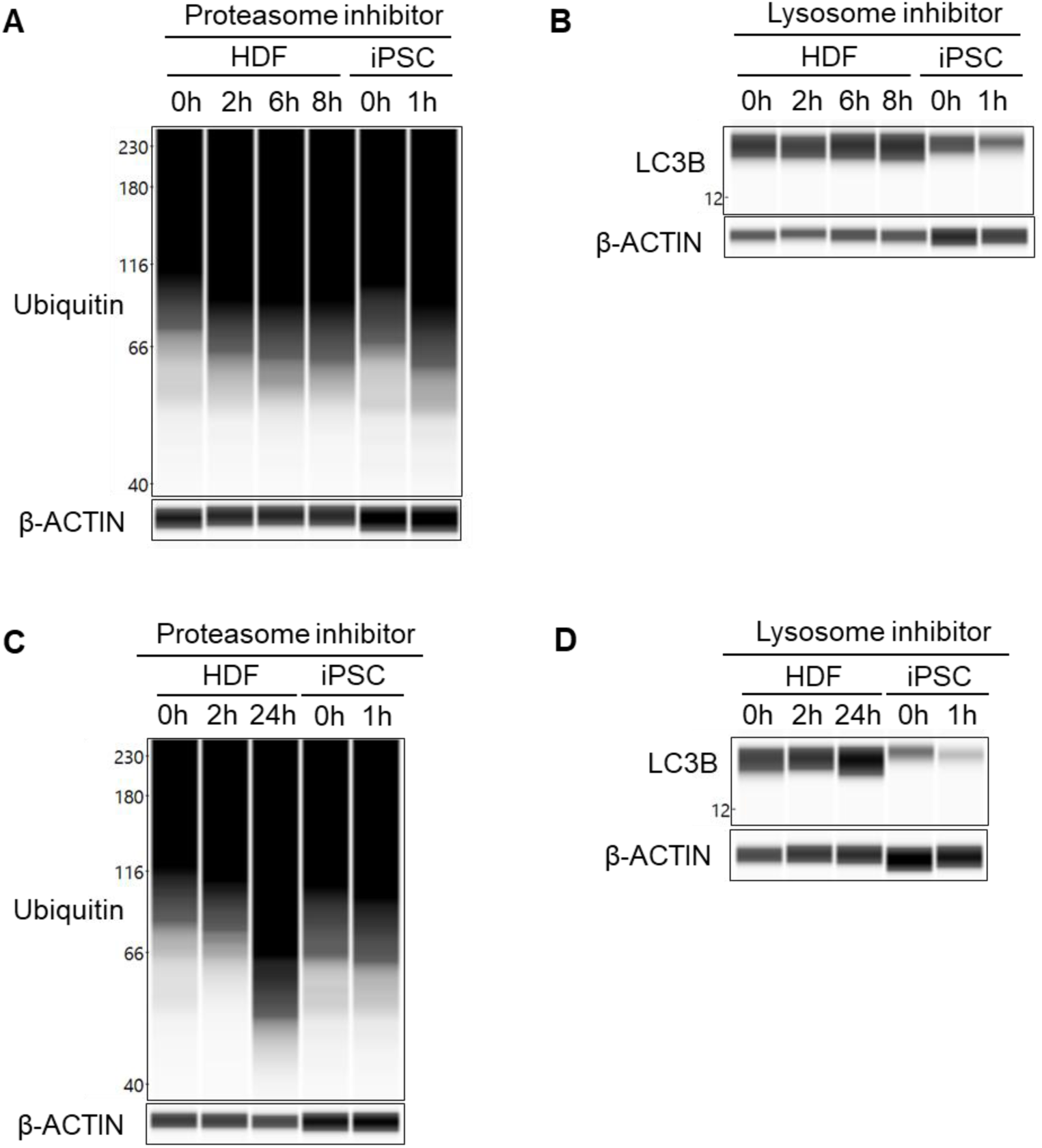
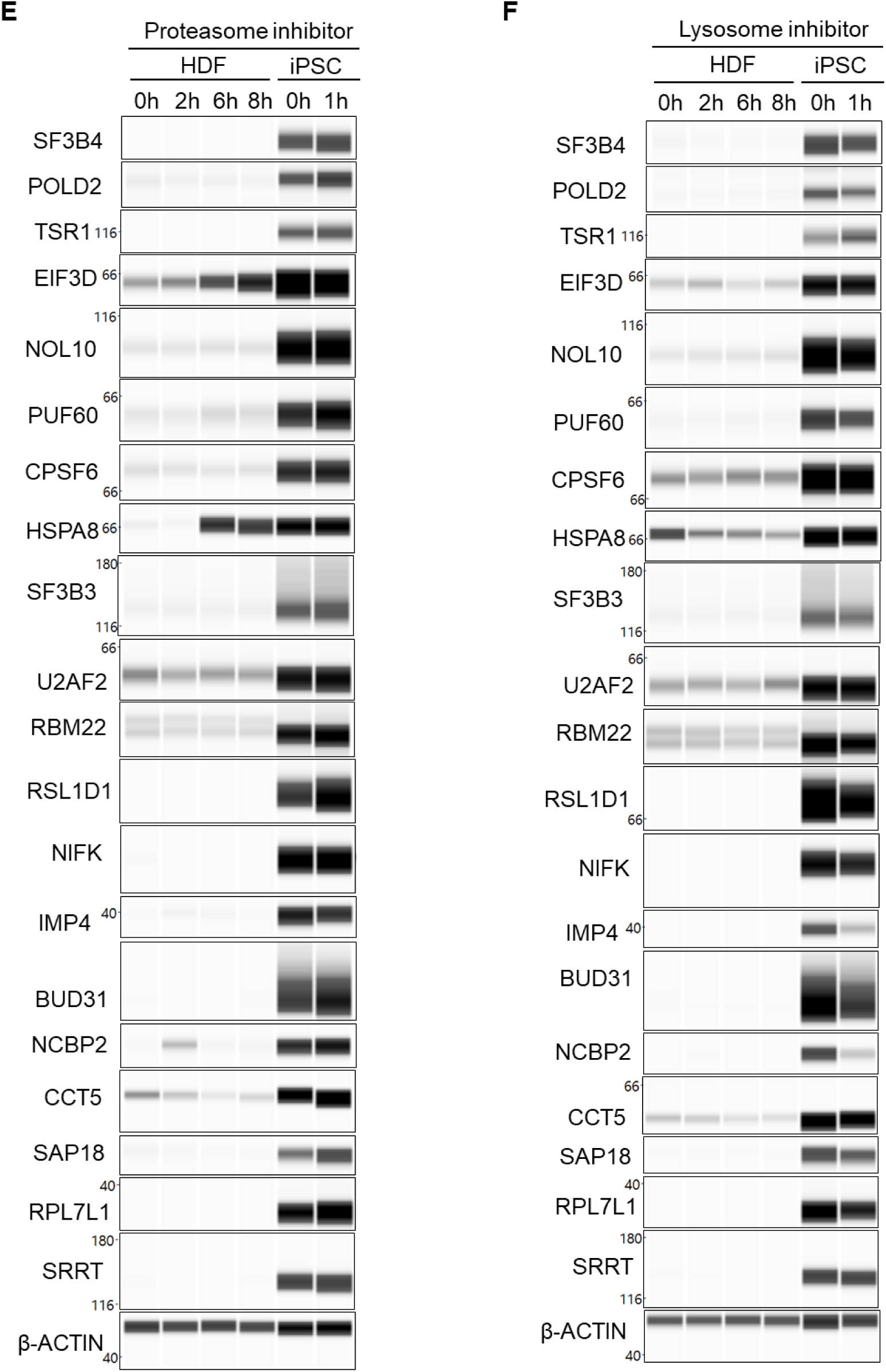
Immunoblotting images of the proteasome inhibitor and lysosomal inhibitor assays. A. Ubiquitin protein levels measured by immunoblotting in iPSC-1 (201B7) and HDF-2 (Tig120) samples treated with a proteasome inhibitor (20 □M MG-132) for up to 8 hours as a control for the proteasome inhibitor assay. B. LC3B protein levels measured by immunoblotting in iPSC-1 (201B7) and HDF-2 (Tig120) samples treated with lysosomal inhibitors (250 nM Bafilomycin A1 and 500 nM Wortmannin) for up to 8 hours as a control for the lysosome inhibitor assay. C. iPSC-1 (201B7) and HDF-2 (Tig120) were treated with 20 □M MG-132 for up to 24 hours, and ubiquitin protein levels were measured by immunoblotting. D. iPSC-1 (201B7) and HDF-2 (Tig120) were treated with 250 nM Bafilomycin A1 and 500 nM Wortmannin for up to 24 hours, and LC3B protein levels were measured by immunoblotting. E. The gene expression levels for the 20 essential PSC-uPRA genes were measured by immunoblotting with the same samples used in A. F. The gene expression levels for the 20 essential PSC-uPRA genes were measured by immunoblotting with the same samples used in B.

**Figure S5.**
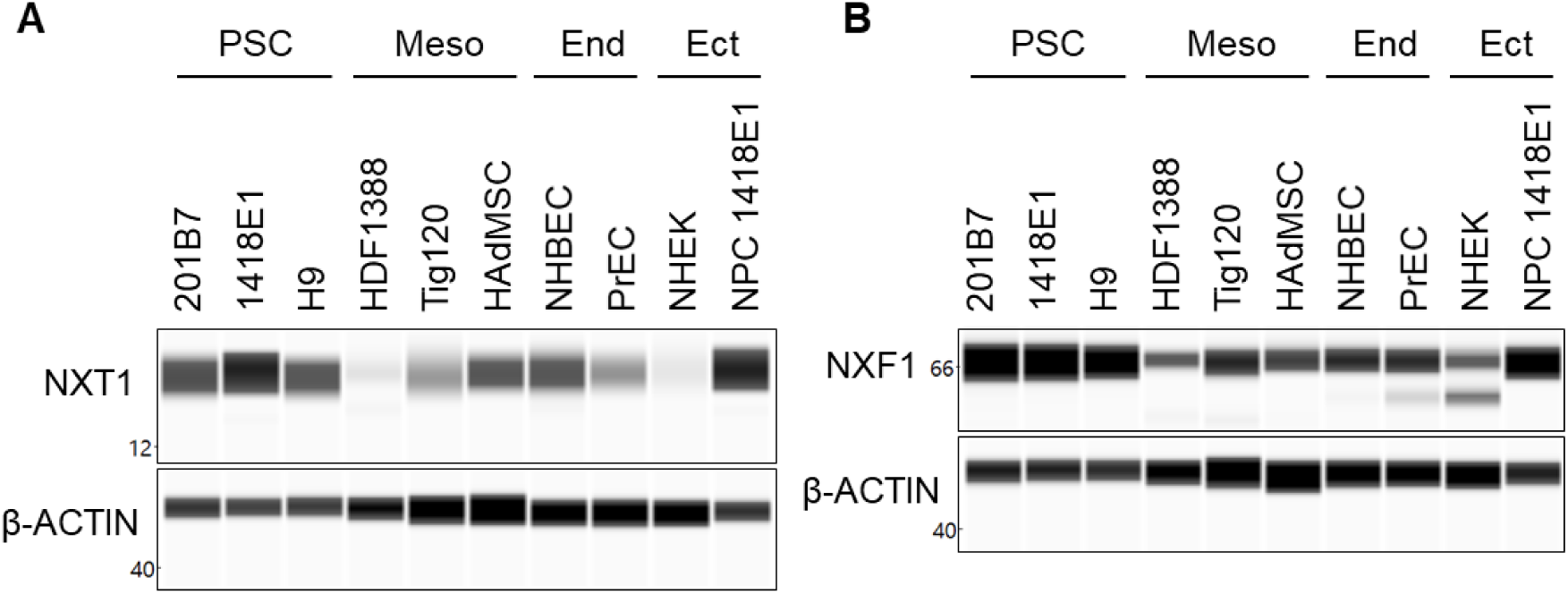
Immunoblotting images of the NXT1 and NXF1 protein expressions between different cell lines Immunoblots of NXT1 (A) and NXF1 (B) for the same samples used in Figure S3 except for NPC 1418E1. Meso, mesoderm; End, endoderm; Ect, ectoderm.

Table S1. Quantified gene list of the mRNA and protein expressions in PSCs and HDFs.

Table S2. List of siRNA used for Figure 2.

Table S3. Number of cells and percentage of immunoassayed proteins following siRNA treatment for 156 targeted genes, related to Figure 2A and B.

Table S4. DNA sequence of the transfected full length FLAG-SRRT.

## ACKNOWLEDGMENTS

We would like to thank Y. Fujita and T. Matsushita for the technical assistance, P. Karagiannis for editorial assistance, and S.Takeshima for administrative support. This work was supported by K-CONNEX from the Japan Science and Technology Agency (JST), Core Center for iPS Cell Research from Japan Agency for Medical Research and Development (AMED; JP21bm0104001), AMED-PRIME (21gm6410003h0001 (M.I.)), and the Japanese Society for the Promotion of Science KAKENHI Grant (JSPS; 19K16104 (M.I.)).

## AUTHOR CONTRIBUTIONS

M.I. and Y.K. performed most of the experiments. T.T generated the program to quantify the MS data. M.Narita performed the microarray. M.Nakagawa., Y.N. and A.O. helped perform the siRNA screening analysis. T.Y. and H.S. performed the mRNA-related experiments. M.I., K.T. and S.Y. designed and directed the research. M.I. analyzed the data and wrote the manuscript with editing by all authors.

