## Supplementary material for "Post-transcriptionally regulated genes are essential for pluripotent stem cell survival": Methods

### KEY RESOURCES TABLE

| REAGENT or RESOURCE | SOURCE | IDENTIFIER |
|---|---|---|
| **Antibodies (dilution ratio used in this study)** | | |
| Rabbit polyclonal anti-BUD31 (1:100) | Proteintech | Cat# 11798-1-AP<br>RRID:AB_2274894 |
| Rabbit monoclonal anti-CCT5 (TCP1e) (1:500) | Abcam | Cat# ab129016<br>RRID:AB_11154964 |
| Rabbit polyclonal anti-CPSF6 (1:100) | Proteintech | Cat# 15489-1-AP<br>RRID:AB_10694140 |
| Rabbit polyclonal anti-EIF3D (1:250) | Proteintech | Cat# 10219-1-AP<br>RRID:AB_2096880 |
| Rabbit monoclonal anti-HSPA8 (D12F2) (1:200) | Cell Signaling Technology | Cat# 8444<br>RRID:AB_10831837 |
| Rabbit polyclonal anti-IMP4 (1:500) | Proteintech | Cat# 16205-1-AP<br>RRID:AB_2239858 |
| Rabbit polyclonal anti-NCBP2 (CBP20) (1:250) | Proteintech | Cat# 11950-1-AP<br>RRID:AB_2150142 |
| Rabbit polyclonal anti-NIFK (MKI67IP) (1:500) | Proteintech | Cat# 12615-1-AP<br>RRID:AB_2142384 |
| Rabbit monoclonal anti-NOL10 (1:500) | Abcam | Cat# ab181161 |
| Rabbit polyclonal anti-POLD2 (1:20) | Proteintech | Cat# 10288-1-AP<br>RRID:AB_2284041 |
| Goat polyclonal anti-PUF60 (1:200) | Abcam | Cat# ab22819<br>RRID:AB_777559 |
| Rabbit polyclonal anti-RBM22 (1:100) | Abcam | Cat# ab157105 |
| Rabbit polyclonal anti-RPL7L1 (1:100) | Proteintech | Cat# 16707-1-AP<br>RRID:AB_1851609 |
| Rabbit monoclonal anti-RSL1D1 (1:2000) | Abcam | Cat# ab181100 |
| Rabbit polyclonal anti-SAP18 (1:100) | Proteintech | Cat# 13841-1-AP<br>RRID:AB_2301353 |
| Rabbit polyclonal anti-SF3B3 (1:1000) | Proteintech | Cat# 14577-1-AP<br>RRID:AB_2270189 |
| Rabbit polyclonal anti-SF3B4 (1:100) | Proteintech | Cat# 10482-1-AP<br>RRID:AB_2301639 |
| Rabbit polyclonal anti-SRRT (ARS2) (1:100) | Abcam | Cat# ab220991 |
| Rabbit polyclonal anti-TSR1 (1:500) | Proteintech | Cat# 16887-1-AP<br>RRID:AB_2222589 |
| Rabbit polyclonal anti-U2AF65 (U2AF2) (1:50) | Proteintech | Cat# 15624-1-AP<br>RRID:AB_2211330 |
| Rabbit monoclonal anti-GAPDH (14C10) (1:500) | Cell Signaling Technology | Cat# 2118<br>RRID:AB_561053 |
| Mouse monoclonal anti-beta-actin (1:250) | Sigma-Aldrich | Cat# A5441<br>RRID:AB_476744 |
| Rabbit polyclonal anti-AFP (1:1000) | Proteintech | Cat# 14550-1-AP<br>RRID:AB_2223933 |
| Rabbit polyclonal anti-MYO1E (1:50) | Proteintech | Cat# 17768-1-AP<br>RRID:AB_2251045 |
| Mouse monoclonal anti-FLAG (M2) (1:40) | Sigma-Aldrich | Cat# F1804<br>RRID:AB_262044 |
| Rabbit polyclonal anti-Ubiquitin (1:50) | Abcam | Cat# ab19247<br>RRID:AB_444805 |
| Rabbit polyclonal anti-LC3B (1:10) | Novus Bio | Cat# NB100-2220SS<br>RRID:AB_791015 |

| | | |
|---|---|---|
| Mouse monoclonal anti-GFP (GF200) (1:100) | Nacalai Tesque | Cat# 04363-24 |
| Mouse monoclonal anti-OCT3/4 (1:500) | BD Biosciences | Cat# BD611202 |
| Rabbit monoclonal anti-NXF1 (1:20) | Abcam | Cat# ab129160<br>RRID:AB_11142853 |
| Mouse monoclonal anti-NXT1 (1:20) | Proteintech | Cat# 67680-1-Ig<br>RRID:AB_2882873 |
| Mouse polyclonal Alexa488 (1:300) | Thermo Fisher Scientific | Cat# A11001<br>RRID:AB_2534069 |
| Rabbit polyclonal Cyanine3 (1:300) | Thermo Fisher Scientific | Cat# A10520<br>RRID:AB_2534029 |
| Rabbit monoclonal anti-LaminB2 (D8P3U) (1:500) | Cell Signaling Technology | Cat# 12255<br>RRID:AB_2797859 |
| Goat anti-rabbit secondary HRP-conjugated (1:1) | Protein Simple | Cat# 042-206<br>RRID:AB_2860577 |
| Goat anti-mouse secondary HRP-conjugated (1:1) | Protein Simple | Cat# 042-205<br>RRID:AB_2860576 |
| Donkey anti-goat secondary HRP-conjugated (1:1) | Protein Simple | Cat# 043-491-2 |
| Chemicals, peptides, Media and recombinant proteins | | |
| SDB-XC Empore disc cartridge | 3M | Cat# 2340 |
| Sodium dodecyl sulfate (SDS) | Nacalai Tesque | Cat# 31606-75 |
| Sodium deoxycholate (SDC) | WAKO | Cat# 190-08313 |
| Sodium lauroyl sarcosinate (SLS) | WAKO | Cat# 192-10382 |
| Sucrose | WAKO | Cat# 195-07925 |
| Lys-C, Mass Spec Grade | Promega | Cat# VA1170 |
| Sequencing Grade Modified Trypsin (Lyophilized) | Promega | Cat# V5111 |
| Ethyl acetate | WAKO | Cat# 051-00356 |
| Acetonitrile | WAKO | Cat# 018-19853 |
| Acetic acid | WAKO | Cat# 018-20061 |
| Methanol | WAKO | Cat# 134-14523 |
| Trifluoroacetic acid (TFA) | WAKO | Cat# 204-02743 |
| Dithiothreitol (DTT) | WAKO | Cat# 045-08974 |
| Iodoacetamide (IAA) | WAKO | Cat# 095-02151 |
| Ammonium bicarbonate | WAKO | Cat# 018-21742 |
| Dimethyl sulfoxide (DMSO) | WAKO | Cat# 045-28335 |
| Triethylammonium bicarbonate buffer | Sigma-Aldrich | Cat# T7408-100ML |
| 1M Tris-HCl (pH 9.0) | Nippon gene | Cat# 314-90381 |
| 1M Tris-HCl (pH 8.0) | Nippon gene | Cat# 314-90065 |
| 1M Tris-HCl (pH 7.5) | Nippon gene | Cat# 316-90221 |
| NaCl (5 M), RNase-free | Thermo Fisher Scientific | Cat# AM9760G |
| $MgCl_2$ (1 M) | Thermo Fisher Scientific | Cat# AM9530G |
| Triton X-100 10% | Teknova | Cat# T1105 |
| Ultrapure water | Kanto Chemical | Cat# 11307-79 |
| Nuclease-Free Water (not DEPC-Treated) | Thermo Fisher Scientific | Cat# AM9937 |
| Y-27632 | Sigma-Aldrich | Cat# Y0503 |
| TrypLE select | Thermo Fisher Scientific | Cat# 12563011 |
| Trypsin-EDTA (0.25%), phenol red | Thermo Fisher Scientific | Cat# 25200056 |

| | | |
|---|---|---|
| Laminin-511 E8 (iMatrix-511) | Nippi | Cat# 892012 |
| StemFiT AK03N | Ajinomoto | Cat# AK03N |
| STEMdiff Neural Progenitor Medium | Stem Cell Technologies | Cat# 05833 |
| Dulbecco's Modified Eagle Medium (DMEM) | Nacalai Tesque | Cat# 08459-35 |
| Fetal Bovine Serum, qualified, New Zealand | Thermo Fisher Scientific | Cat# 10091148 |
| BEGM Bronchial Epithelial Cell Growth Medium BulletKit | Lonza | Cat# CC-3170 |
| PrEGM Prostate Epithelial Cell Growth Medium BulletKit | Lonza | Cat# CC-3166 |
| MesenPRO RS Medium | Thermo Fisher Scientific | Cat# 12746012 |
| STEMdiff Neural Progenitor Medium | STEMCELL Technologies | Cat# ST-05833 |
| ReagentPack Subculture Reagents | Lonza | Cat# CC-5034 |
| InSolution MG-132 | Calbiochem | Cat# 474791 |
| Bafilomycin A1 | InvivoGen | Cat# tlrl-baf1 |
| InSolution WORTMANNIN | Calbiochem | Cat# 681676 |
| TurboDNase (2U/uL) | Thermo Fisher Scientific | Cat# AM2238 |
| SUPERaseIn | Thermo Fisher Scientific | Cat# AM2694 |
| Cycloheximide solution, 100 mg/mL in DMSO | Sigma-Aldrich | Cat# C4859-1ML |
| Protease Inhibitor | Sigma-Aldrich | Cat# P8340-1ML |
| cOmplete Mini, EDTA-free Protease Inhibitor Cocktail | Sigma-Aldrich | Cat# 11836170001 |
| Phosphatase Inhibitor cocktail 2 | Sigma-Aldrich | Cat# P5726-1ML |
| Phosphatase Inhibitor cocktail 3 | Sigma-Aldrich | Cat# P0044-1ML |
| Critical commercial assays | | |
| SurePrint G3 Human GE 8x60K v3 | Agilent Technologies | Cat# G4858A |
| iTRAQ Reagents Multiplex Kit | Sciex | Cat# 4352135 |
| Subcellular Protein Fractionation Kit for Cultured Cells | Thermo Fisher Scientific | Cat# 78840 |
| BCA Protein Assay Kit | Thermo Fisher Scientific | Cat# 23225 |
| Quantitative Fluorometric Peptide Assay | Thermo Fisher Scientific | Cat# 23290 |
| Anti-Rabbit Detection Module for Jess, Wes, Peggy Sue or Sally Sue | Protein Simple | Cat# DM-001 |
| Anti-Mouse Detection Module for Jess, Wes, Peggy Sue or Sally Sue | Protein Simple | Cat# DM-002 |
| 12-230 kDa Jess or Wes Separation Module, 8 x 25 capillary cartridges | Protein Simple | Cat# SM-W004 |
| miRNeasy Mini Kit | QIAGEN | Cat# 217004 |
| Cellstain - Hoechst 33342 solution | Dojindo | Cat# 346-07951 |
| QIAzol lysis reagent | QIAGEN | Cat# 79306 |
| Trizol LS reagent | Thermo Fisher Scientific | Cat# 10296028 |
| SuperScript III First-Strand Synthesis SuperMix for qRT-PCR | Thermo Fisher Scientific | Cat# 11752050 |
| ReverTraAce | TOYOBO | Cat# TRT-101 |
| In-Fusion HD Cloning Kit | Clontech Laboratories | Cat# 639648 |
| TaqMan Gene Expression Master Mix | Thermo Fisher Scientific | Cat# 4369016 |

| | | |
|---|---|---|
| PowerUp SYBR Green Master Mix | Thermo Fisher Scientific | Cat# A25742 |
| Amicon Ultra centrifugal filters (10K) | Merck Millipore | Cat# UFC501096 |
| Lipofectamine Stem Reagent | Thermo Fisher Scientific | Cat# STEM00015 |
| Lipofectamine MessengerMAX | Thermo Fisher Scientific | Cat# LMRNA001 |
| Opti-MEM Reduced Serum Medium | Thermo Fisher Scientific | Cat# 31985062 |
| Stemfect RNA Transfection Kit | STEMGENT | Cat# 00-0069 |
| Deposited data | | |
| Raw and analyzed data (Human gene expression microarray) | This study | GSE184546 |
| Raw images | Mendeley | DOI: 10.17632/y6b3bgng9p.1 |
| Raw and analyzed data (mass spectrometry) | jPOST | JPST001308 (PXD028489) |
| Experimental models: Cell lines | | |
| 201B7 human iPSC line | (Takahashi et al., 2007) | RRID:CVCL_A324 |
| 1418E1 human iPSC line | This study | N/A |
| HDF1388 Human dermal fibroblast | Purchased from Cell applications, Inc. | N/A |
| Tig120 Human dermal fibroblast | National Institutes of Biomedical Innovation, Health and Nutrition. | N/A |
| H9 embryonic stem cell | (Thomson et al., 1998) | WA09 RRID:CVCL_9773 |
| Normal human bronchial epithelial cells; NHBEC | Lonza Bioscience | Cat# CC-2541 |
| Human prostate epithelial cells; PrEC | Lonza Bioscience | Cat# CC-2555 |
| Human adipose tissue-derived mesenchymal stem cells; HAdMSC | Life Technologies | Cat# R7788115 |
| Normal human epidermal keratinocytes; NHEK | Lonza | Cat# 00192627 |
| H9 ESC-derived neural progenitor cells | Thermo Fisher Scientific | N7800200, RRID:CVCL_IU37 |
| 1418E1 iPSC-derived neural progenitor cells | This study | N/A |
| Oligonucleotides | | |
| BUD31 (Hs00696974_m1) TaqMan Assay | Thermo Fisher Scientific | Cat# 4331182 |
| CCT5 (Hs04362335_g1) TaqMan Assay | Thermo Fisher Scientific | Cat# 4331182 |
| CPSF6 (Hs01101212_m1) TaqMan Assay | Thermo Fisher Scientific | Cat# 4331182 |
| EIF3D (Hs01044815_m1) TaqMan Assay | Thermo Fisher Scientific | Cat# 4331182 |
| HSPA8 (Hs03044880_gH) TaqMan Assay | Thermo Fisher Scientific | Cat# 4331182 |
| IMP4 (Hs00369187_m1) TaqMan Assay | Thermo Fisher Scientific | Cat# 4331182 |
| NCBP2 (Hs01597558_g1) TaqMan Assay | Thermo Fisher Scientific | Cat# 4331182 |
| NIFK (Hs00757500_s1) TaqMan Assay | Thermo Fisher Scientific | Cat# 4331182 |

| | | |
|---|---|---|
| NOL10 (Hs01042161_m1) TaqMan Assay | Thermo Fisher Scientific | Cat# 4331182 |
| POLD2 (Hs00371757_g1) TaqMan Assay | Thermo Fisher Scientific | Cat# 4331182 |
| PUF60 (Hs01050525_g1) TaqMan Assay | Thermo Fisher Scientific | Cat# 4331182 |
| RBM22 (Hs00216159_m1) TaqMan Assay | Thermo Fisher Scientific | Cat# 4331182 |
| RPL7L1 (Hs02339924_g1) TaqMan Assay | Thermo Fisher Scientific | Cat# 4331182 |
| RSL1D1 (Hs00378363_g1) TaqMan Assay | Thermo Fisher Scientific | Cat# 4331182 |
| SAP18 (Hs00705532_s1) TaqMan Assay | Thermo Fisher Scientific | Cat# 4331182 |
| SF3B3 (Hs00418633_m1) TaqMan Assay | Thermo Fisher Scientific | Cat# 4331182 |
| SF3B4 (Hs00538859_g1) TaqMan Assay | Thermo Fisher Scientific | Cat# 4331182 |
| SRRT (Hs00210818_m1) TaqMan Assay | Thermo Fisher Scientific | Cat# 4331182 |
| TSR1 (Hs00250762_m1) TaqMan Assay | Thermo Fisher Scientific | Cat# 4331182 |
| U2AF2 (Hs00200737_m1) TaqMan Assay | Thermo Fisher Scientific | Cat# 4331182 |
| ACTB (Hs01060665_g1) TaqMan Assay | Thermo Fisher Scientific | Cat# 4331182 |
| GAPDH (Hs02786624_g1) TaqMan Assay | Thermo Fisher Scientific | Cat# 4331182 |
| 18S (Hs99999901_s1) TaqMan Assay | Thermo Fisher Scientific | Cat# 4331182 |
| MALAT1 (Hs00273907_s1) TaqMan Assay | Thermo Fisher Scientific | Cat# 4331182 |
| GammaTub23C (Dm01841764_g1) TaqMan Assay | Thermo Fisher Scientific | Cat# 4331182 |
| siRNAs, see Table S2 | Dharmacon | N/A |
| DNA sequence for mRNA transfection, see Table S4 | This study | N/A |
| **Software and algorithms** | | |
| Excel 2016 | Microsoft | https://www.office.com/ |
| ProteinPilot v5.0 | Sciex | https://sciex.com/products/software/proteinpilot-software |
| Mascot | Matrix Science | https://www.matrixscience.com/ |
| Compass | Protein Simple | https://www.proteinsimple.com/compass/downloads/ |
| HCS Studio Cell Analysis Software | Thermo Fisher Scientific | N/A |
| GeneSpring version 14.9.1 | https://www.agilent.com/ | Agilent Technologies |
| TargetMine | Mizuguchi Laboratory https://doi.org/10.3389/fgene.2019.00934 | https://targetmine.mizuguchilab.org/targetmine/ |

| | | |
|---|---|---|
| Removal of interference mixture MS/MS spectra (RiMS) perl script | Mio Iwasaki Laboratory (Iwasaki et al., 2019) | https://sourceforge.net/projects/rimsprogram/files/ |
| Other | | |
| MonoCap C18 HighResolution 4000 (0.1 x 4000 mm) | GL Sciences | Cat# 5020-4000 |

### CONTACT FOR REAGENT AND RESOURCE SHARING

### METHOD DETAILS

#### Cell culture

Two types of human induced pluripotent stem cells (iPSC-1, 201B7; and iPSC-2, 1418E1) and embryonic stem cells (ESC; H9) were maintained under feeder-free conditions on iMatrix-511 (Nippi) with StemFit AK03N medium (Ajinomoto). Karyotypes of both iPSCs were verified as normal. A Rock inhibitor (Y-27632, final concentration, 10 µM) was used only on the day of plating (Nakagawa et al., 2014). Two types of human dermal fibroblasts (HDF-1, HDF1388; and HDF-2, Tig120) were cultured in Dulbecco's modified Eagle's medium (DMEM; Nacalai Tesque) with 10% fetal bovine serum (FBS, Thermo Fisher Scientific) and 0.5% penicillin and streptomycin (gibco). Endodermal cells (normal human bronchial epithelial cells, NHBEC; and human prostate epithelial cells, PrEC), mesodermal cells (human adipose tissue-derived mesenchymal stem cells, HAdMSC), and ectodermal cells (normal human epidermal keratinocytes, NHEK) were cultured according to the manufacturer's harvesting protocol (Lonza). Human neural progenitor cells derived from H9 (NPC H9) and 1418E1 (NPC 1418E1) were cultured in STEMdiff Neural Progenitor Medium according to the manufacturer's harvesting protocol (Veritas).

#### Protein extraction

The cells were washed once with ice-cold PBS (Nacalai Tesque) and directly lysed with ice-cold lysis buffer (PTS buffer: 12 mM SDC, 12 mM SLS, 100 mM Tris-HCl (pH 9.0)) or SDS buffer (1% SDS, 20 mM Tris-HCl (pH 8.0)), with 1% phosphatase and protease inhibitors (Sigma-Aldrich). Cell lysates were collected by scraping and pipetting on pre-chilled Protein LoBind 2 mL tubes (Eppendorf). After sonication and heat shock at 95$^{o}$C for 5 min, the protein concentration was determined using a BCA protein assay kit (Thermo Fisher Scientific, #23227).

#### Gene expression analysis by capillary-based immunoblotting assay

The immunoblotting analysis was performed using a Wes automated capillary electrophoresis system (Protein Simple) according to the manufacturer's protocol. The protein concentration was aligned to around 0.5 mg/mL of the sample. Information about the primary and secondary antibodies and the dilutions are provided in KEY RESOURCES TABLE. The data was analyzed and visualized using Compass for Simple Western software (Protein Simple).

**Gene expression analysis by qRT-PCR and microarray**

The cells were lysed with QIAzol lysis reagent (QIAGEN) for whole cell lysis, and fractionated samples were lysed with TRIZOL LS reagent (Thermo Fisher Scientific). Total RNA was purified using a miRNeasy Mini Kit (QIAGEN). Purified RNA (0.1-1 µg) was used for single strand complementary DNA (cDNA) synthesis using a SuperScript III First-Strand Synthesis SuperMix for qRT-PCR (Thermo Fisher Scientific) or ReverTra Ace (TOYOBO). Quantitative RT-PCR was performed using TaqMan Gene Expression Master Mix (Thermo Fisher Scientific) or PowerUp SYBR Green Master Mix (Thermo Fisher Scientific) on a Quant Studio 3 instrument (Thermo Fisher Scientific). The mRNA levels were normalized to the human GAPDH or drosophila gamma-Tubulin at 23C (gammaTub23C) expression, and then the relative expressions were normalized with the control. Microarray was performed as described previously (Takahashi et al., 2020) using the purified total RNA.

**Subcellular fractionation of mRNA and proteins**

Subcellular fractionations of cytoplasm, organelles, and nuclei were performed using the Subcellular Protein Fractionation Kit for Cultured Cells (Thermo Fisher Scientific, #78840). Briefly, the cells were washed once with ice-cold PBS (Nacalai Tesque). After completely removing PBS, the cells were lysed with 200 µL (24-well plate), 500 µL (6-well plate), or 1000 µL (9-cm dish) of CEB solution at 4ºC for 5 min. Cell lysates were collected by scraping and pipetting on pre-chilled Protein LoBind 2 mL tubes (Eppendorf). The lysates were centrifuged at 500 x g for 5 min, and then the supernatants were collected as cytoplasmic fractions. The cell pellets were dissolved using 200 µL (24-well plate), 500 µL (6-well plate), or 1000 µL (9-cm dish) of MEB solution. After a 5-s vortex, the lysate tubes were inverted for 10 min at 4ºC. The supernatants were collected as organelle fractions after centrifugation at 3000 x g for 5 min. The pellets were collected as nucleus fractions. For protein extraction, the cytoplasm and organelle fractions were ultra-filtrated using an AmiconUltra-0.5 (10K) at 14000 x g for 10 min and solubilized by PTS buffer. For the nucleus fractions, 200 µL (24-well plate), 500 µL (6-well plate), or 1000 µL (9-cm dish) of PTS buffer was added to solubilize the pellet. Protein extraction was performed as described above. For RNA extraction, an equal volume of TRIZOL LS reagent (Thermo Fisher Scientific) was added to the cytosol and organelle fractions. For the nucleus fraction, 400 µL (24-well plate) or 1000 µL (6-well plate) of QIAzol lysis reagent (QIAGEN) was added to solubilize the pellet. RNA extraction was performed as described above.

**Gene expression analysis by nanoliquid chromatography (nanoLC)-mass spectrometry (MS)**

Protein samples lysed with PTS buffer were subjected to reduction, alkylation, Lys-C/trypsin digestion (enzyme ratio 1/100), and desalting as previously described (Iwasaki *et al.*, 2019). The resulting peptides were labeled with isobaric tags for relative and absolute quantification (Sciex). Briefly, 120 µg of desalted peptide samples were dried and dissolved in 10 µL of 500 mM triethyammonium bicarbonate. Approximately 20 µL of iTRAQ reagents (Sciex) was added to 23 µL of ethanol and mixed with the peptide sample. After incubation for 1.5 h at room temperature, 16 µL of 10% TFA and 400 µL of loading buffer (0.5% trifluoroacetic acid and 4% (v/v) acetonitrile) were added to quench the reaction, and the sample mixture was desalted using StageTip (Rappsilber et al., 2003). A total of 8 µg of iTRAQ labelled sample set was subjected to nanoLC-MS/MS using a TripleTOF 5600 System (AB Sciex) equipped with an HTC-PAL autosampler (CTC Analytics). All nanoLC-MS conditions were the same as previously described (Iwasaki *et al.*, 2019) including the monolithic column (4 m length, 100 µm i.d., GL Science) and total analytical time of 1080 min. NanoLC-MS/MS analysis was performed in triplicate, and blank runs were inserted between the samples. The proteome data analysis method was previously reported (Iwasaki *et al.*, 2019). For quantification, the accumulated intensity of iTRAQ label spectra was calculated after the application of the RiMS method to remove the interference spectra (Iwasaki *et al.*, 2019). Then, the final normalized accumulated intensity of iTRAQ label spectra was calculated for the whole and subcellular fraction samples in technical triplicates to acquire the protein quantification value for each cell type.

**Trans-omics data analysis**

The mRNA and protein ratio between cell types was calculated using the normalized mRNA and protein quantification values described above for iPSC-1, iPSC-2, H9, HDF-1, and HDF-2. Then, the mRNA and protein ratios of PSC/HDF were compared based on the same gene symbol or Entrez Gene. To determine statistical significance, we conducted unpaired t tests with Excel 2016 (Microsoft) in biological triplicate. The definition of independent mRNA and protein upregulation is as follows. Gene ontology analysis was performed using TargetMine (Chen et al., 2019).

mRNA-Up with no protein change: -0.9 < Protein ratio < 0.9 (Log2), mRNA ratio >= 0.9 (Log2)

mRNA-Down with no protein change: -0.9 < Protein ratio < 0.9 (Log2), mRNA ratio <= -0.9 (Log2)

Protein-Up with no mRNA change: -0.9 < mRNA ratio < 0.9 (Log2), Protein ratio >=0.9 (Log2)

Protein-Down with no mRNA change: -0.9 < mRNA ratio < 0.9 (Log2), Protein ratio <= -0.9 (Log2)

**siRNA screening**

201B7 and 1418E1 were seeded on iMatrix-511-coated 96-well plates at 4,940 cells/well. Y-27632 (final concentration, 10 µM) was used for the iPSC maintenance for two days after passage and transfection. Tig120 was seeded directly on 96-well plates at 2,470 cells/well. On the next day of the passage (day 1), a manually aliquoted siGENOME SMARTpool siRNA library (0.2 µM, 7.5 µL/well; Horizon Discovery) was mixed with an equal volume of 0.5% of Stemfect RNA Transfection Reagent (STEMGENT). Then, 20 µL of siRNA solution and 80 µL of fresh medium were mixed, and the cell medium was replaced with 80 µL of this solution. On the next day, the medium of the wells with transfection reagent was manually replaced. As controls for the siRNA screening, we used siNontarget, siOCT3/4, and LaminB2 for every 3 wells per plate. Four days after the transfection, cells were fixed and analyzed by immunostaining. In brief, the cells were washed twice with PBS (Nacalai Tesque) and fixed with 4% paraformaldehyde (PFA) for 10 min at room temperature. Then the fixed cells were treated with PBS containing 0.2% Triton X-100 and 1% bovine serum albumin (BSA, Thermo Fisher Scientific) for 15 min at room temperature after two PBS washes. The cells were incubated with the primary antibody anti-OCT3/4 (B&D) or Lamin B2 (CST) diluted in PBS containing 1% BSA for more than one hour at room temperature with protection from light. After washing twice with PBS, the cells were incubated with 0.01% Hoechst 33342 (Dojindo) and the secondary antibody Alexa488 or Cy3 diluted in PBS containing 1% BSA for one hour at room temperature with protection from the light. After two PBS washes, PBS was added to the well, and 10x immunofluorescence images were acquired using an ArrayScan VTI 600 Series (Thermo Fisher Scientific). Obtained images were analyzed using HCS Studio (Thermo Fisher Scientific, version 6.5.0, build 7614). The siRNA knockdown of OCT3/4 and LaminB2 was respectively used as a pluripotency marker for PSCs and a universal control for both cell types (PSCs and HDFs). We measured the cell number by Hoechst staining and checked the general knockdown efficiency by immunostaining. The average knockdown efficiencies by siOCT3/4 were 100.0% and 99.7% for 201B7 (iPSC-1) and 1418E1 (iPSC-2), respectively, and by siLaminB2 it was 98.4% for Tig120 (HDF-2). The criteria for a change in cell numbers by siRNA against OCT3/4, LaminB2, or no target was a value more than ±2SD the mean cell number and checked manually. Dilution conditions of the primary and secondary antibodies are provided in KEY RESOURCES TABLE. A list of the siRNA used in this study is provided in Table S2.

#### Proteasome and lysosome inhibitor assay

One semiconfluent 6-well dish of adherent 201B7 or Tig120 was treated with medium containing 20 µM MG-132 (Calbiochem) or 250 nM Bafilomycin A1 (InvivoGen) and 500 nM Wortmannin (Calbiochem) for the proteasome or lysosome assays, respectively. After 0, 2, 6, 8, and 24 h, the cells were washed once with ice-cold PBS (Nacalai Tesque) and directly lysed with ice-cold PTS buffer with 1% phosphatase and protease inhibitors (Sigma-Aldrich). Cell lysates were collected by scraping and pipetting on pre-chilled Protein LoBind 2 mL tubes (Eppendof). After sonication and heat shock at 95$^{o}$C for 5 min, the protein concentration of the sample was determined using the BCA protein assay kit.

#### Monosome and polysome fractionation

One semiconfluent 100-mm dish of adherent 201B7 or Tig120 was treated with medium containing 100 µg/mL Cycloheximide (Sigma-Aldrich) for 5 min at 37$^{o}$C. The cells were placed on ice and gently washed once with 10 mL ice-cold PBS. Then they were lysed with 0.4 mL ice-cold lysis buffer (20 mM Tris-HCl, pH7.5, 150 mM NaCl, 5 mM $MgCl_2$, 1 mM dithiothreitol (DTT), Complete EDTA-free Protease Inhibitor Cocktail, 100 µg/mL Cycloheximide, 1% Triton X-100, 25 units/mL Turbo DNase (Thermo Fisher Scientific), and 100 units/mL SUPERaseIn (Thermo Fisher Scientific)), scraped, and collected into a 1.5 mL chilled DNA LoBind Tube (Eppendorf). The lysate was incubated on ice for 10 min and triturated through a 25-gauge needle (Terumo) ten times before centrifugation at 20,000 xg for 10 min at 4$^{o}$C. The supernatant was collected in a new 1.5 mL tube for the sucrose gradient analysis and as the sample for loading. A 10%–45% continuous sucrose gradient was prepared in a polyclear tube (Seton) using 10% and 45% sucrose buffers containing 100 µg/mL Cycloheximide and 1 mM DTT in polysome buffer (25 mM Tris-HCl (pH 7.5), 150 mM NaCl and 15 mM $MgCl_2$) and the Biocomp Gradient Master program (Biocomp). An equal amount of cell lysate as sample (300 mL) was loaded on the prepared gradient solution. Monosome and polysomes were separated in the sucrose gradient by ultracentrifugation using a SW-41 rotor (Beckman Coulter) at 36,000 rpm for 2.5 h at 4$^{o}$C. The profile of relative RNA abundances of monosomes and polysomes were visualized at 254-nm wavelength, and equal-volume fractions were collected simultaneously with the Biocomp Piston Gradient Fractionator (Biocomp). For the RNA analysis, an equal sample volume of TRIzol LS reagent (Thermo Fisher Scientific) was immediately added to the fractions and load sample. RNA was purified using an miRNeasy Mini Kit according to manufacturer's instruction. Purified RNAs along with 1 ng of spiked drosophila RNA were used for the cDNA synthesis and the following qRT-PCR. The cycle threshold for detectable gene expression was set as Ct = 45. Ct values were normalized by the spiked drosophila RNA and compared with the loading sample before the sucrose gradient.

### mRNA transfection

One day before the transfection, we plated 201B7 and Tig120 at a density of 2 x $10^5$ cells and 0.7 x $10^5$ cells per well on a 24-well plate, respectively. For the iPSCs, we prepared an LN511-coated 24-well plate, and Y-27632 (final concentration, 10 µM) was used for the iPSC maintenance for two days after passage. For the mRNA transfection, N1-methyl-pseudouridine (1mΨ)-modified mRNA was prepared, and 100 ng mRNA was transfected using Lipofectamine MessengerMAX (Thermo Fisher Scientific) on the next day of passage (day 1) according to the manufacturer's protocol. The next day, the cells were washed twice with PBS (Nacalai Tesque) and lysed with 150 µL PTS buffer.

### DATA AVAILABILITY

Gene expression microarray results are accessible in the Gene Expression Omnibus (GEO) database of the National Center for Biotechnology Information website (accession number: GSE184546). The mass spectrometry data have been deposited to the ProteomeXchange Consortium via jPOSTrepo (Okuda et al., 2017) (https://repository. jpostdb.org/) with the data set identifier JPST001308 (PXD028489) for the 2-plex analysis of iPSC-1 (201B7) and HDF-1 (HDF1388), and for the 4-plex analysis of iPSC-1 (201B7), iPSC-2 (1418E1), ESC (H9), and HDF-2 (Tig120). Original immunoblot data have been deposited to Mendeley Data (DOI: 10.17632/y6b3bgng9p.1).
